# Positive interactions are common among culturable bacteria

**DOI:** 10.1101/2020.06.24.169474

**Authors:** Jared Kehe, Anthony Ortiz, Anthony Kulesa, Jeff Gore, Paul C. Blainey, Jonathan Friedman

## Abstract

Interspecies interactions shape the structure and function of microbial communities. In particular, positive, growth-promoting interactions can significantly affect the diversity and productivity of natural and engineered communities. However, the prevalence of positive interactions and the conditions in which they occur are not well understood. To address this knowledge gap, we used kChip, an ultra-high throughput coculture platform, to measure 180,408 interactions among 20 soil bacteria across 40 carbon environments. We find that positive interactions, often described to be rare, occur commonly, primarily as parasitisms between strains that differ in their carbon consumption profiles. Notably, non-growing strains are almost always promoted by strongly growing strains (85%), suggesting a simple positive interaction-mediated approach for cultivation, microbiome engineering, and microbial consortium design.

**One Sentence Summary:** Experimental measurement of >150,000 bacterial cocultures reveals that growth-promoting interactions occur commonly and depend on differences in nutrient consumption preferences.

## Main Text

Microbial communities are composed of multiple species that interact with one another in a variety of ways as part of the “struggle for existence” (*1*). Interactions between species are broadly classified as either negative interactions, where a species inhibits another species’ growth through nutrient exploitation and chemical warfare (*2*); or positive interactions, where a species promotes another species’ growth by increasing nutrient availability and creating new niches (*3*). These positive interactions can also be bi-directional, forming a mutualism in which two species mutually aid each other’s growth. The overall distribution of positive and negative interactions within a microbial community profoundly impacts the community’s structure, stability and productivity (*4*–*6*). These properties, in turn, shape a community’s ability to perform vital functions for the environment (*7*–*10*) and for host organisms (*11*–*14*). Despite the importance of the distribution of microbial interactions, the relative prevalence of positive and negative interactions in nature remains largely unknown.

Positive interactions are generally thought to be rare. Experimental evidence from coculture studies points to a dominance of negative interactions (*2, 15*). For example, evidence of positive interactions was found in <10% of pairs of bacteria isolated from tree holes (*15*). However, these results are subject to strong experimental biases, such as the use of a single environment—even though microbial interactions can differ dramatically across environmental conditions (*16*–*18*)—and the use of strains that each grow individually in the environmental conditions being tested. Metabolic modeling, which can simulate millions of interactions across myriad environments, as well as limited experimental evidence, suggest that positive interactions emerge via environment-dependent secretions and can facilitate otherwise non-growing species (*18*–*21*). Additionally, evolutionary theories like the Black Queen Hypothesis argue that such secretion-mediated positive interactions are selected for (*22*). Taken together, these findings suggest that positive interactions among microbes may be common and play a significant role in shaping microbial communities, but such theories have not yet been thoroughly tested experimentally.

Quantifying the prevalence of positive interactions and determining the conditions in which they occur could significantly improve our ability to predict and control the ecology of microbial communities (*23, 24*). Positive interactions are predicted to enhance a community’s diversity and productivity but decrease its stability (*6, 25, 26*). Therefore, a better understanding of these interactions would enhance our ability to manipulate and manage communities, with widespread applicability in environmental conservation (*27*), crop health (*28*), and human health (*29*). Nevertheless, the data required for quantifying the distribution of interactions across environments is still lacking due to methodological limitations that frustrate comprehensive sampling of interactions under many conditions (*30*). Inferring interactions from metagenomic sequencing remains an outstanding challenge (*31, 32*), and directly measuring interactions at scale is difficult to perform using existing experimental paradigms.

To gain a broad understanding of how species interact across a wide range of environments, we used a combinatorial screening platform called kChip (*33*–*35*) to measure >150,000 pairwise bacterial cocultures among 20 different soil bacterial strains across 40 environments with differing carbon source identity or concentration. The kChip generates cocultures at an ultra-high throughput scale by rapidly and randomly combining droplets containing microbial cultures and/or medium components within microwells (*33*) (**Fig. 1A-D, fig S1**). Here, we paired unlabeled (wild-type) and GFP-labeled versions of 20 Gammaproteobacteria isolated from soil (6 *Enterobacterales* and 14 *Pseudomonadales*, **Table S1, fig S2 and S3**). We selected these bacterial strains from our larger pool of fluorescently labeled soil isolates maximizing for phylogenetic diversity (**Table S2, Materials and Methods**). We cocultured each strain pair in each of 33 single carbon sources (0.5% w/v), a mix of these, five of the 33 at reduced concentration (0.05% w/v), and a no-carbon control. Carbon sources were drawn from several biochemical classes including carbohydrates, amino acids, and carboxylates (**Table S3**). We measured the effect of each unlabeled strain on the growth of each labeled strain in each carbon source, giving a total of 17,600 combinations (20 labeled strains, [20 + 2 control] unlabeled strains, and [39 + 1 control] carbon sources), each represented >10 times on average in our data (**fig S4**). These one-way interactions were used to classify each pairwise interaction qualitatively (**Fig 1E**) and quantitatively (**Fig 1F**).

**Fig. 1.**
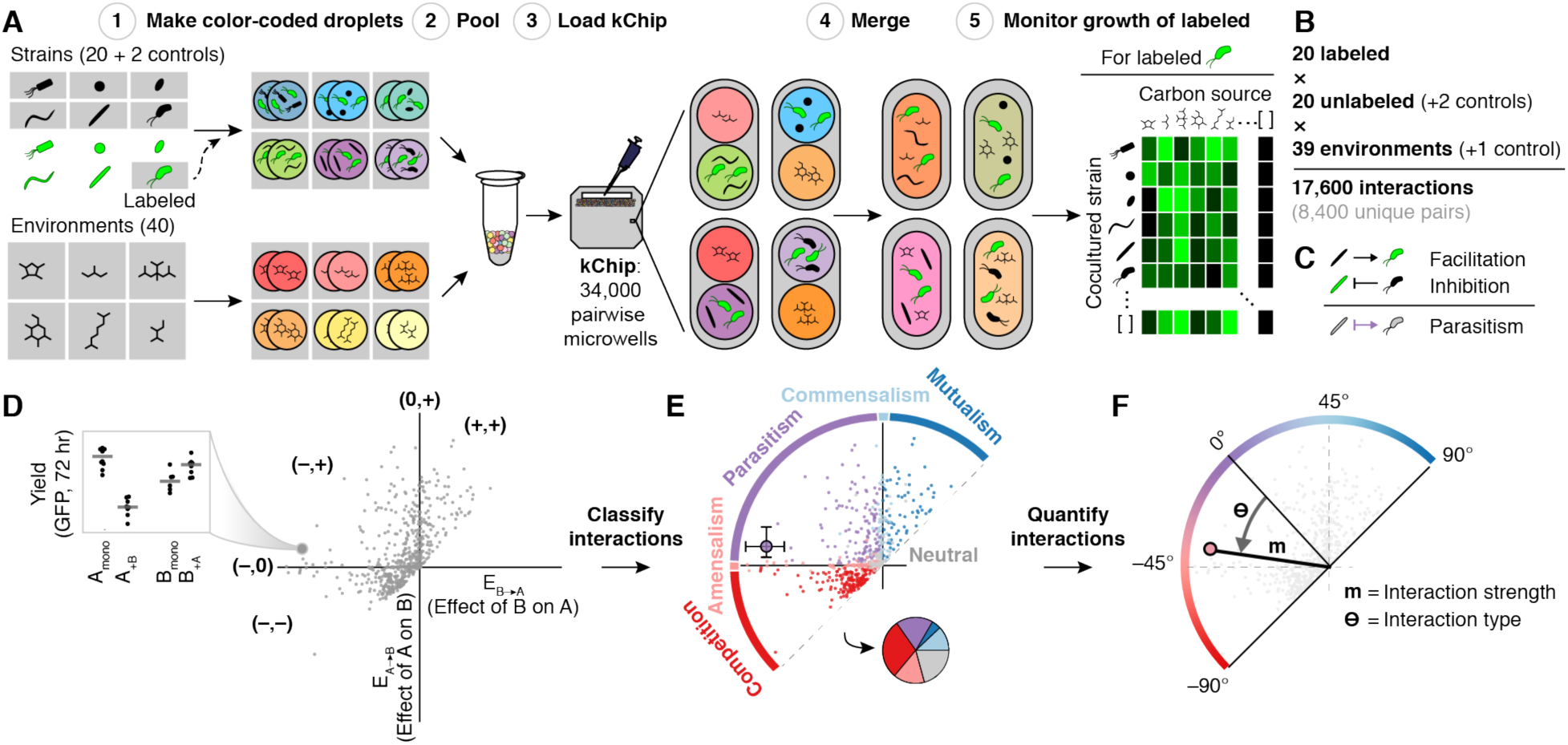
High-throughput interaction assay and analysis. **A.** Steps to assay the effect of multiple unlabeled species on a single label species across carbon sources on each kChip. Color-coded droplets, each containing either a labeled + unlabeled coculture or a single carbon source, were generated (Step 1), pooled together (Step 2) and loaded onto a kChip (Step 3). Each kChip contained an array of microwells that randomly paired coculture droplets with carbon source droplets. After imaging the color codes to identify the inputs per microwell, droplet pairs were merged via exposure to an electric field (Step 4), and the growth of the labeled strain was measured at 0, 24, and 72 hrs (Step 5). **B.** Overall size of the kChip screen. **C.** Using data across kChips, bidirectional interactions were deduced by combining data where each strain within a given pair was the labeled strain. **D-F.** Framework for kChip data analysis. Each pairwise interaction was described qualitatively (interaction classification) and quantitatively (interaction strength, m and interaction type, Θ).

Our data provide direct experimental evidence that positive interactions are indeed common, and occur primarily as parasitisms. More broadly, we found that interactions strongly depend on the environment via differences in the carbon consumption preferences of the interacting strains. Notably, we discovered that strongly growing partners consistently enabled the growth of strains that were unable to grow in monoculture (85%), suggesting a simple strategy for cultivation, microbiome engineering, and design of microbial consortia.

### Positive interactions occur frequently

The effect of each unlabeled strain on each labeled strain was classified as positive (yield increase compared to monoculture), negative (yield decrease compared to monoculture), or 0 if there was no evidence for an effect (**Fig 1C, D, S5**) (see **Materials and Methods**). Bidirectional pairwise interactions were classified as mutualism (+,+), commensalism (0,+), parasitism (–,+), amensalism (–,0), competition (–,–), or neutralism (0,0) (**Fig 1E, fig S6**). We focused our analysis on the 72 hr time point when cultures had reached saturation.

Positive interactions were common overall (**Fig 2A**). Excluding cases when neither strain within a pair grew detectably in monoculture, >40% of cocultures contained ≥1 positive interaction, with over half of these occurring within a parasitism (22% parasitisms, 14% commensalisms, and 5% mutualisms). There were also many cocultures that contained only negative interactions (35% competitions and 18% amensalisms). Relatively few interactions were neutralisms (6%), though neither strain grew in monoculture for 21% of pairs.

**Fig. 2.**
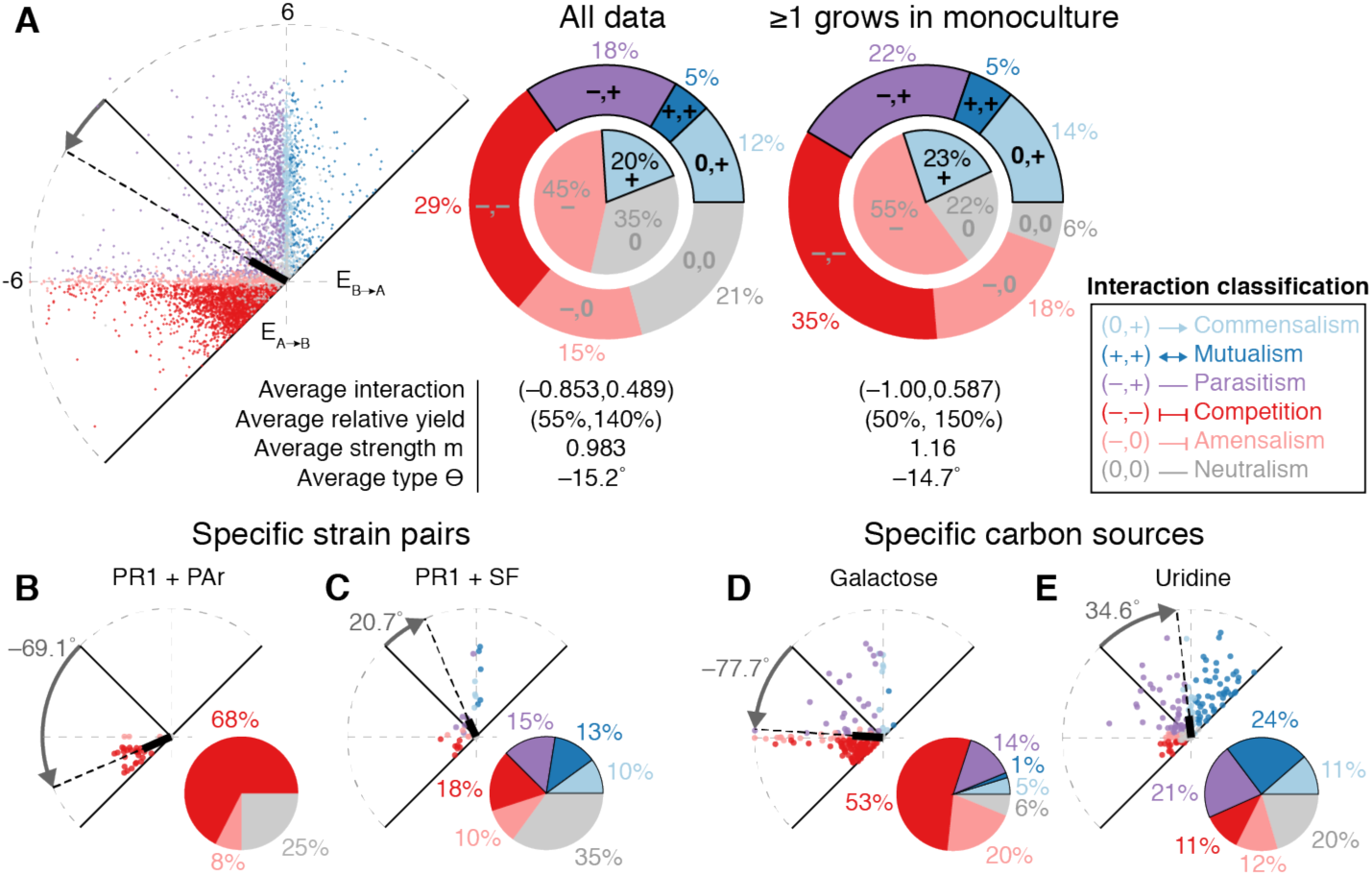
Positive interactions occurred commonly and depended on strain pair and carbon source. **A.** (Left) All pairwise interactions on 33 distinct carbon sources. (Middle) Interaction classification of all data. The inner circle represents one-way interactions classifications; the outer ring represents two-way interaction classifications. (Right) Interaction classification excluding cases in which both strains comprising a coculture showed no detectable growth as monocultures on a given carbon source. **B-C.** Interactions of two example cocultures on all carbon sources. **D-E.** Interactions of all cocultures on two example carbon sources. Colors indicate interaction classification (legend). All data at 72 hr.

The extent to which interacting pairs influenced each other’s growth also varied quantitatively across strain and carbon environments. For example, in a parasitism, the facilitated strain may have increased in yield more than the inhibited strain decreased or vice versa. To capture these quantitative differences, we measured the magnitude m (strength) and angle Θ (type) of each pairwise interaction in polar coordinates (**Fig 1F**). Θ represented the relative effect of cocultured strains on each other (with Θ = −90° indicating that both strains inhibited each other equally, Θ = 0° indicating a balanced parasitism with equal and opposite effects, and Θ = 90° indicating both strains facilitated each other equally).

The average interaction was a parasitism (Θ = −14.7° and m = 1.16) in which the inhibited strain was hurt (2X decrease) more than the facilitated strain was helped (1.5X increase) (**Fig 2A**). The results at 24 hr were similar and included even more positive interactions, with 54% of cocultures containing ≥1 positive interaction (20% parasitisms, 25% commensalisms, and 9% mutualisms), although the average interaction, also a parasitism, favored the facilitated strain (1.3X decrease and 1.7X increase) (**fig S7**).

### The occurrence of positive interactions differs among strain pairs and among carbon sources

The prevalence of positive interactions differed significantly among strain pairs: Positive interactions never occurred for some pairs, while for others they occurred in a large fraction of the tested environments (**Fig 2B-C**). Many pairs were capable of producing all six interaction types across the carbon source library (**Fig 2C**), indicating that it can be challenging to predict the interaction of a particular pair in a given environment based on the pair’s interaction in a different environment.

Positive interactions were also more common on certain carbon sources than others. In galactose, for example, a majority of pairwise interactions were competitions (53%), with only 20% containing ≥1 positive interaction (**Fig 2D**). In uridine, however, competitions were relatively rare (11%), with 56% of pairwise interactions containing ≥1 positive interaction (**Fig 2E**). Several carbon sources were capable of producing all interaction classes across our strain set (**fig S8**).

The strong dependence of the interaction type on strain identity and carbon source indicated a need to examine our dataset broadly in order to detect patterns governing the occurrence of positive interactions. We therefore tested whether characteristics of the environmental conditions or the strain pairs could explain the interactions we observed.

### Positive interactions occurred independently of environmental characteristics

Broad environmental characteristics, including carbon source biochemical class and number of carbon atoms, did not appear to dictate the occurrence of positive interactions. We found that the variation in interaction classification between carbon sources within a class was not significantly different than variation between classes (**Fig 3A**) (pairwise PERMANOVA, *p* > 0.05). We hierarchically clustered all interactions for all strain pairs and carbon sources and found that carbon sources did not group by biochemical class (TCA cycle carboxylate ions were the singular exception) (**fig S9**). We further found that the occurrence of positive interactions was largely independent of the number of carbon atoms each carbon source possessed (**Fig 3B**) (Spearman correlation = 0.32, *p* = 0.07). For example, serine, a compound with only 3 carbon atoms, enabled more positive interactions than all other carbon sources except uridine.

**Fig. 3.**
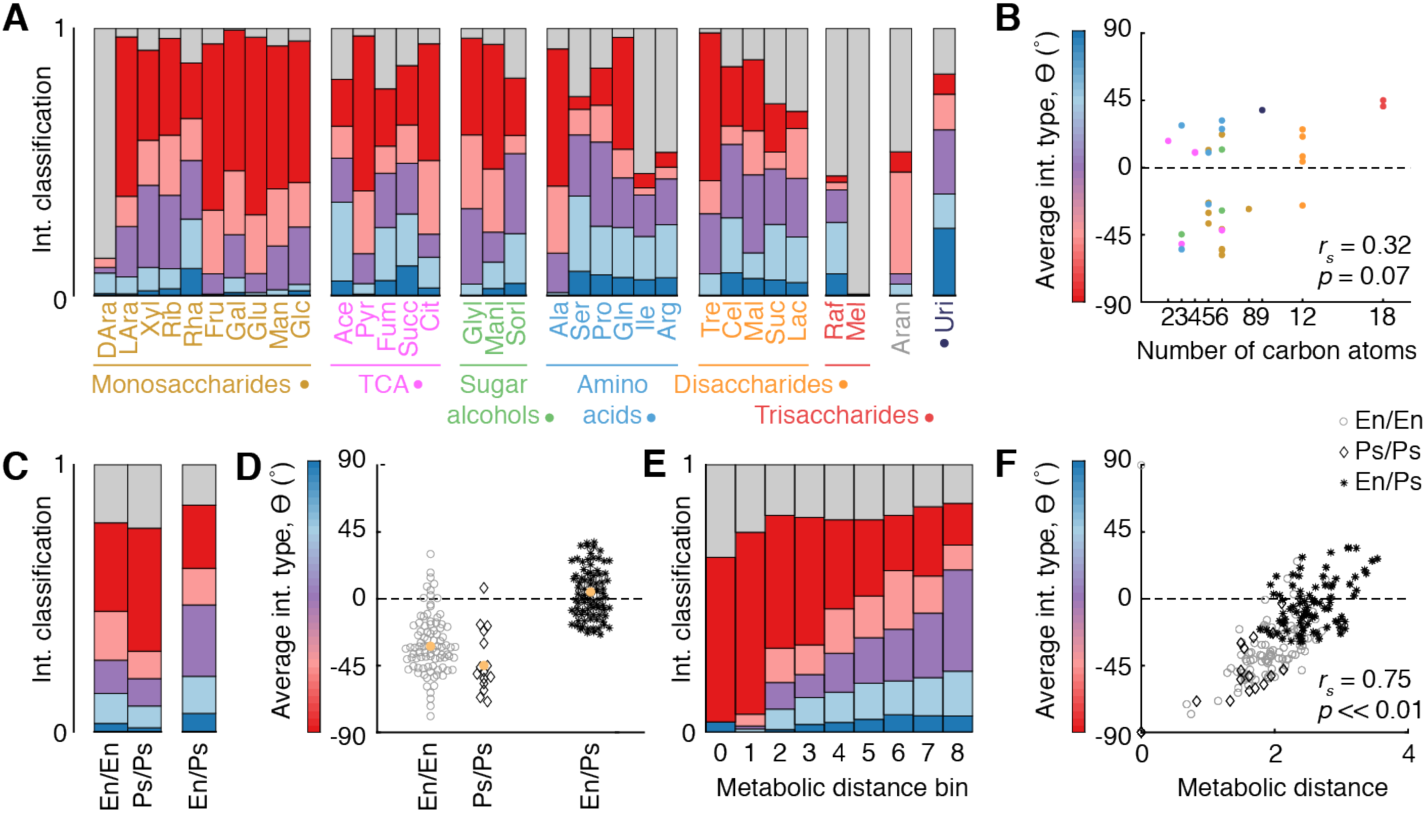
Positive interactions depended strongly on strain properties. **A.** Interaction classification by carbon source organized into biochemical categories. Bar colors indicate interaction classification. **B.** Average interaction type by number of carbon atoms. Dot color indicates biochemical class. **C.** Interaction classification by phylogenetic relatedness of the taxonomic pairs. En = *Enterobacterales*. Ps = *Pseudomonadales*. **D.** Interaction type by phylogenetic relatedness of the taxonomic pairs. **E.** Interaction classification by pairwise Euclidean metabolic distance (binned). Bin 0 represents with-self interactions. Bins 1-8 each contain roughly equal numbers of pairwise interactions. **F.** Interaction type by metabolic distance. All data at 72 hr.

### Positive interactions increase with strain dissimilarity

We found that positive interactions were more common among taxonomically dissimilar strains. We compared interactions within and between the two taxonomic orders represented in our set, *Enterobacterales* (En) and *Pseudomonadales* (Ps), since pairwise phylogenetic distances, calculated from full-length 16S rRNA gene sequences (**fig S3**), distributed bimodally (**fig S10**). More positive interactions occurred among inter-order pairs (En+Ps) than intra-order pairs (En+En and Ps+Ps) (**Fig 3C**): 26% of En+En pairs and 20% of Ps+Ps pairs contained ≥1 positive interaction, compared to 47% for En+Ps pairs. The average interaction type was negative for intra-order cocultures (−32° for En+En and −45° for Ps+Ps) but positive for inter-order cocultures (4.5° for En+Ps) (**Fig 3D**) without significant differences in interaction strength (**fig S11**).

The fraction of positive interactions, especially parasitisms, increased with the metabolic distance between the interacting species (**Fig 3E**). Metabolic differences between strains were calculated as the Euclidean distances between strains’ carbon source utilization profiles, given by each strain’s ability to grow on each carbon source in monoculture (**fig S2**). The distribution of metabolic distances between all strains was a continuous bell-shaped distribution (**fig S10**), which captured more finely graded functional differences among strains than bimodal phylogenetic distances. Intra-strain interactions (i.e. those between labeled strains and their unlabeled counterparts) were typically competitions (62% of all strains, and 90% when excluding non-growing strains) that reduced the yield of the labeled strain by 50% on average (**fig S12**). Intra-strain mutualisms occurred in a small fraction of strains (3.7%), which we hypothesize was caused by higher starting density (a phenomenon known as an Allee effect (*36*)). Interactions between metabolically similar but non-identical strains were also typically competitions (68%) (**Fig 3E**). As metabolic distance increased, the fraction of pairs that exhibited ≥1 positive interaction increased monotonically from 0.2% to 61%, with a growing fraction of these positive interactions occurring within parasitisms. This result indicated a strong and directional dependence of positive interaction frequency on the functional dissimilarity of strain pairs.

As metabolic distance increased, the average interaction type Θ also became increasingly positive (Spearman correlation = 0.75, *p* << 0.01) (**Fig 3F**). This trend reflects both shifts in the frequency of interaction classes and quantitative changes in the relative strength of interactions’ components. Notably, among parasitisms, the average Θ within each metabolic distance bin increased, indicating an increasingly large facilitative effect relative to the reciprocal inhibitory effect (**fig S11**). Since the prevalence of interaction types was correlated with strains’ metabolic capabilities, we next tested whether interactions in a given environment were dependent on the interacting strains’ abilities to grow in monoculture in that environment.

### Monoculture yields shape occurrence of positive interactions in coculture

Interaction networks for each carbon source suggested that monoculture yield indeed related to interactions (examples in **Fig 4A** and all networks in **fig S13**). Regardless of the carbon sources or strain pair, positive interactions (often in the form of parasitisms) commonly occurred between strong and weak growers and sometimes occurred between weak growers; competition commonly occurred between strong growers (**Fig 4B**).

**Fig. 4.**
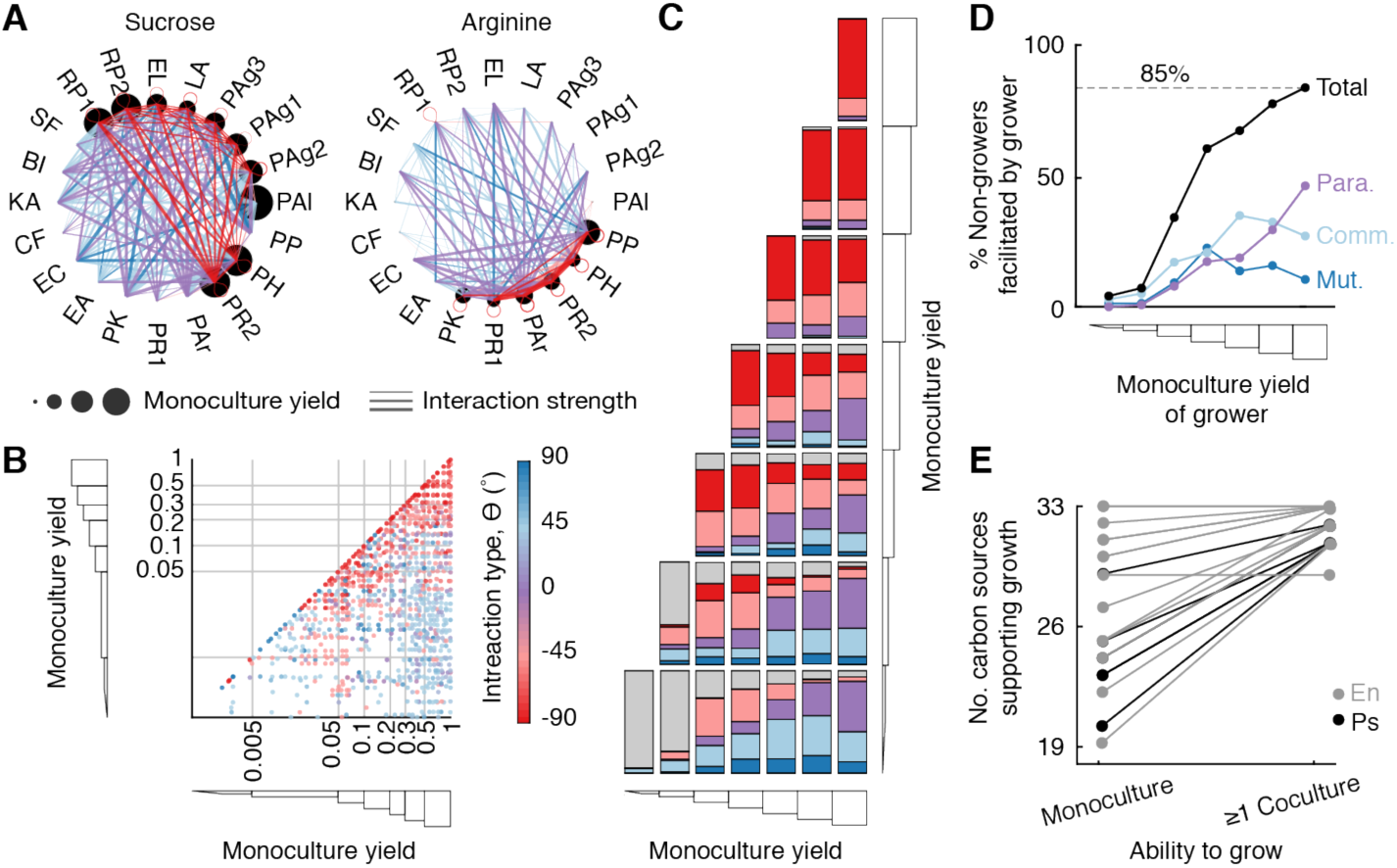
Carbon source utilization capabilities shape interactions. **A.** Example interaction networks for two carbon sources, each with different subsets of strains that could utilize it. Node size represents strain growth in monoculture (background-subtracted and normalized to each strain’s maximum monoculture yield). Edge color represents interaction classification. Edge thickness represents interaction strength (m) **B.** Interaction types (color) by the monocultures yields of both strains in each coculture. Gray lines indicate monoculture growth bins. **C.** Interaction classification for each pair of monoculture growth bins. **D.** Fractions of non-growers that are obligately facilitated by strains with different monoculture yields. Line colors represent the interaction classifications in which the facilitation occurs (Mut. = mutualism; Comm. = commensalism; Para. = parasitism; Total = total fraction of non-growers facilitated). **E.** The total number of carbon sources on which each strain can grow in monoculture and in at least one coculture. Each line represents one strain. All data at 72 hr.

These patterns were found to be robust in a systematic analysis of interactions as a function of monoculture yield (**Fig 4B-C**). As monoculture yield became more dissimilar, the frequencies of mutualisms, commensalisms, and parasitisms all increased (off-diagonal of **Fig 4C** and **fig S14B**). By contrast, for interactions of strain pairs able to grow on a carbon source equally well (diagonal of **Fig 4C** and **fig S14A**), competitions were consistently prevalent and increased in frequency with monoculture yield. Among interactions between the strongest growers, 77% were competitions where both species were inhibited compared to their monoculture growth. Moreover, in 99% of pairs of the strongest growers at least one species was inhibited (77% competitions, 18% amensalisms, and 4% parasitisms). Interaction at 24 hr followed the same trend (**fig S15**), with the overall increased prevalence of positive interactions (**fig S7**) likely reflecting the lack of high monoculture yield values. Overall, interactions appeared to depend heavily on the two interacting strains’ individual abilities to grow in a specific environment, and this dependence explained the interaction variability exhibited by certain strain pairs and certain carbon sources.

These results were also consistent with the fact that there were far fewer positive interactions on low-concentration carbon sources (11% with ≥1 positive interaction, as opposed to 35% for their high-concentration counterparts) (**fig S16**). The low-concentration carbon sources supported low-to-mid-range monoculture yields that were similar among the strains (**fig S2**). There were few instances of strong growers paired with weak growers, the regime where positive interactions otherwise emerged (**fig S16**). Consequently, the interaction classifications on low-concentration carbon sources were nearly as different to their high-concentration counterparts as the high-concentration carbon sources were to each other (**fig S17**). Unlike these low-concentration carbon sources, a mix of 33 carbon sources produced consistently high monoculture yields (most similar to glucose) (**fig S1**). As a result, interactions occurred only between strong growers and were consequently highly negative (85% competitions, with 99% containing ≥1 inhibition) (**fig S18**). Indeed, for each strain pair, the interaction type Θ on the mixed carbon source condition was, with a single exception (PR1+PAr), always lower than its average interaction across all carbon sources (**fig S18**). Altogether, differences in monoculture yields provided a common explanation for differences in the interaction distributions across timepoints, carbon source types, and carbon source concentrations.

### Non-growing strains are typically facilitated by strongly growing strains

To quantify the prevalence of obligate facilitation, where a non-growing strain required a facilitating partner to grow on a given carbon source, we examined interactions between strains that could grow strongly on a carbon source and those that could not (bottom row of **Fig 4C**, and **fig S14C**). As the yield of the growing strain increased, the frequency of facilitation of the non-grower increased markedly. When paired with the strongest growers, non-growers were facilitated 85% of the time. Moreover, the relative fraction of these positive interactions that occurred as parasitisms also increased (**Fig 4D**), making up 55% of obligate facilitations between the strongest growers and non-growers. Among these parasitisms, the average interaction type also became increasingly positive (**fig S15**): The magnitude of the positive interaction increased monotonically without the inhibited strain experiencing the same degree of yield loss (**fig S19**).

Of the 20 strains, we found that all but one (PAl) were capable of growing in coculture with a facilitating strain on any carbon source they could not utilize in monoculture (**fig S19**). Of the 125 instances where a strain did not grow on a carbon source (6.25 carbon sources per strain on average), only 21 persisted across all cocultures (1.05 carbon sources on average) (**Fig 4E, fig S20**). That is, independent of the carbon source and strain identity of the non-grower, at least one of the 19 candidate partner strains were typically observed that enabled growth. This common obligate facilitation suggests how biodiversity can be supported when few carbon sources are available or when only a subset of strains can utilize available carbon sources.

## Discussion

In this study, we performed comprehensive pairwise coculturing of 20 culturable soil microbes across 40 different carbon source environments. By measuring interactions across many environments, our study produced many instances in which at least one of the two cocultured strains grew poorly or not at all, a regime in which we found positive interactions were far more likely to occur than previously measured. The study also produced instances where both strains grew well and antagonism was common, a result consistent with previous large-scale studies of bacterial interactions (*37, 38*).

Our study unearthed a wealth of positive interactions. While mutualisms were relatively rare (5%), commensalisms (12%) and parasitisms (18%) were common, and accounted for the majority of cases where total coculture yield was greater than the sum of monocultures (24%) (**fig S21**)—a criterion previously used to classify cooperative interactions (*15*). The prevalence of these positive interactions corroborates predictions from large-scale metabolic models (*19, 20, 39*). Our results are also consistent with the predictions of theories like the Black Queen Hypothesis, which asserts that interspecies cross-feeding of “leaky” public goods are evolutionarily selected for (*22, 40*). Finally, our results generalize smaller scale demonstrations that cocultured strains (*41*) and spent media (*42*) can induce growth of fastidious bacteria. Altogether, positive interactions increasingly appear to play a dominant role in driving community properties, such as resistance to invasion and productivity (*3, 22*), and in supporting microbial biodiversity (*43*).

Interactions varied significantly across environments and timepoints (**fig S19**). This suggests that interactions, and therefore properties of natural communities, can display considerable spatial and temporal variability. While interactions were not significantly dependent on properties intrinsic to the environment itself, they nonetheless strongly depended on the environment via the ability of each strain to individually grow on it: Negative interactions were frequent between strong growers, while positive interactions occurred commonly between strong and weak growers across all timepoints and environments. Therefore, given the widespread differences in growth that occur among bacteria, positive interactions may occur commonly in nature.

A variety of mechanisms could explain the prevalence of positive interactions in our data. First, facilitated strains might have grown on components of accumulating dead cells, though this is unlikely given the timescale of the coculture experiment (*44*). Second, the facilitator might have secreted carbon source-degrading enzymes that increased overall carbon availability. This mechanism is consistent with the general prevalence of positive interactions in dimeric and trimeric sugars (**Fig 3A**) but may not explain positive interactions in simple carbon sources like monomeric sugars and TCA cycle intermediates (**Fig 3A, B**). Third, the facilitator may have excreted incompletely oxidized metabolites that were utilized by the facilitated strain (*20, 40, 45*). Such “overflow metabolism” would allow strains to indirectly benefit from the biochemical transformation capabilities of their facilitators. Exploitation of newly created niches could explain the positive interactions we observed on simple carbon sources (e.g. the excretion of short-chain fatty acids as a byproduct of incomplete monosaccharide oxidation). It may also explain the rarity of positive interactions on lower carbon source concentrations, since respiration is known to be favored over fermentation under such conditions and overflow is less likely to occur (*45*).

Despite the high throughput of our experiment, we were limited in the number of strains and carbon sources relative to the diversity that exists in nature. Our strain library was limited to two taxonomic orders isolated from topsoil. Additionally, we only chose strains that grew on a minimal medium as part of our culturing protocol, possibly biasing our dataset against obligate interactions involving amino acid or vitamin auxotrophies, which are known to be common (*46*). Moreover, while our carbon source library represented a variety of carbon source types, it was limited to soluble compounds, excluding many polymers on which metabolically-driven positive interactions may be more common. Whether our results extend across additional phylogenetic groups and nutrient environments should be investigated in follow-up studies.

Our results indicate that knowledge of how strains grow individually in an environment can be strongly predictive of how they interact in that environment. In contrast, knowing how the same strains interact in a different environment or how different strains interact in the same environment do not appear to be very informative. Finally, our results suggest that a potential strategy for inducing the growth of a non-growing or weakly growing strain, independent of growth medium, is to coculture it with a strongly growing strain. Here we uncovered several general, statistical rules governing microbial community structure and function. Such rules deepen our understanding of microbial community ecology and are crucial to enable the efficient design and control of beneficial microbial communities.

## Acknowledgments

The authors thank Seppe Kuehn and Nadav Kashtan for critical comments on the manuscript; Alfonso Perez Escudero, Otto Cordero, Eric Alm, Pardis Sabeti, and Cheri Ackerman for useful discussions; Megan Hoffman for assistance; and the Jupyter, numpy, scipy, scikit-image, scikit-learn, and pandas open source development teams.

## Author contributions

This work was conceptualized by all authors. The kChip experimental platform was designed by J.K., A.K., and P.C.B. Microbial isolation was performed by J.K. and A.O. Microbial strain labeling was performed by A.O. Software was written by J.K. and A.K. Formal analysis was conducted by J.K. with feedback from A.O., P.C.B., and J.F. The experimental investigation was carried out by J.K. and A.O. Experimental data was curated by J.K. and phylogenetic data was curated by A.O. The original manuscript draft and data visualization was prepared by J.K. The manuscript was reviewed and edited by all authors. The study was supervised by J.G., P.C.B., and J.F.

## Data and materials availability

The kChip platform designs and screen data collected herein are available upon request.

## Supplementary Materials

Materials and Methods

Figures S1-S21

Tables S1-S3

