## Supplement for "Positive interactions are common among culturable bacteria"

**This PDF file includes:**

Materials and Methods

Figs. S1 to S21
Tables S1 to S3

### Materials and Methods

Strain isolation from soil samples

Soil samples (two ~12cm columns of topsoil, ~4cm in diameter) were collected from multiple locations in greater Boston on November 12, 2017 (5.6°C) (specific locations listed in **Table S1**). Each sample was diluted in PBS within a few hours of collection (5 g soil vortexed in 40 mL PBS). Single strains were first isolated by streaking 70 µL of dilutions of this mixture ($10^{-1}$, $10^{-2}$, $10^{-3}$, and $10^{-4}$) on 20 different solid (agar) media [Tryptic Soy Broth (TSB) (Bacto), 1% v/v TSB, Lysogeny Broth (LB) (Bacto), 1% v/v LB, Nutrient Broth (NB) (Bacto), 1% v/v NB, M9 salts (Sigma-Aldrich) + 0.5% w/v glucose, M9 salts + 0.005% w/v glucose, M9 salts + 0.005% w/v glucose + 0.2% w/v casamino acids, M9 salts + 0.005% w/v glucose + 0.002% w/v casamino acids, M9 salts + 0.5% w/v glucose + 0.2% w/v casamino acids at pH = 4 and 5, M9 salts + 0.005% w/v glucose + 0.002% w/v casamino acids at pH = 4 and 5, Actinomycete Isolation Agar (Teknova), Brucella Agar (Teknova), Streptomyces Medium (Teknova), Campylobacter Medium (Teknova), Bordetella Medium (Teknova), and ATCC Medium 1111 (Teknova)].

Strains were selected based on the following criteria: growth in LB liquid medium of transferred colony (25°C), frozen glycerol stock revival in LB ($OD_{600}$ > 0.1) (30°C), and subsequent growth on M9 + 0.5% w/v glucose ($OD_{600}$ > 0.1) (30°C). We kept 96-well plates of isolates (LB, 25% v/v glycerol) at -80°C.

Isolates included in the coculture experiment (**Table S1**) were selected based on robust revival from glycerol stocks and in subsequent culturing steps and the ability to label the strains via constitutive expression of GFP.

Strain labeling

The soil bacterial isolates were fluorescently labeled with the commercially available plasmid pMRE132 containing GFP2 (*47*). First, the soil isolates and the *E. coli* carrying pMRE132 (*Ec*-pMRE) were grown to saturation (5 mL of LB media [Bacto] in 50 mL Falcon tubes, loose caps, 30°C, 24 hrs, 15mg/L Chloramphenicol [Sigma-Aldrich] for *Ec*-pMRE). Second, the saturated culture (SC) of each soil isolate was mixed with the SC of *Ec*-pMRE (500 µL of each). This and subsequent steps were done in 2 mL 96 deep-well plates (Eppendorf) with a Viaflo 96 liquid handler (Integra) to increase throughput. The mixed SCs were concentrated to 10X by centrifuging (1 min, 7,000 RCF), discarding 900 µL of the supernatant, and re-suspending. Immediately after, 10 µL of each SC was spotted, in three replicates, onto nutrient agar (5 g peptone, 3 g yeast extract, 15 g agar [Bacto] in 1L water). The agar plates were incubated (24 hrs, 30°C) to allow for conjugation and transfer of the plasmid from *Ec*-pMRE to the soil isolates. The bacterial lawns were picked with sterile wooden applicators (Puritan), suspended in phosphate buffered saline (Corning), and mixed by pipetting 30 times with Viaflo. The suspensions were then serially diluted ($10^0$, $10^{-1}$ and $10^{-2}$) and spotted onto minimal media agar to select for the transconjugants (1X M9 minimal salts [Teknova], 5 g/L Glucose, 15 mg/L Chloramphenicol [Sigma-Aldrich], 15 g agar [Bacto]). Fluorescent colonies were picked and streaked onto minimal media agar to counterselect *Ec*-pMRE. Finally, SC from single colonies were frozen with 50% glycerol and stored at -80°C.

Strain identification and calculation of phylogenetic distance

Each bacterium isolated from soil was classified phylogenetically with its 16S rRNA gene sequence. The full 16S gene sequences (~1500bp) were obtained via Sanger sequencing (Quintara Biosciences), quality threshold-trimmed, and classified with Seqmatch (*48*). *Sulfolobus* *solfataricus*, a thermophilic archaeon, was used in the phylogenetic reconstruction as an outgroup species to root the tree. MUSCLE (*49*) with default parameters was used to align the sequences. PhyML-SMS (*50*, *51*) with default parameters was used to select GTR+G+I as the best model and to infer the tree. The inferred branching order and branch lengths in the phylogenetic tree are the maximum likelihood estimate given the sequence alignment. The pairwise phylogenetic distances between strains were calculated directly from the phylogenetic tree with R (Phytools).

Media construction

The medium used in the coculture experiment was an M9 minimal medium consisting of 1X M9 salts (Teknova), 1X trace metals (Teknova), 0.1 mM calcium chloride, and 2 mM magnesium sulfate. We additionally added 0.05% w/v bovine serum albumin (BSA) to the medium to improve the retention of fluorescent dyes used in the droplet color codes (*33*) (see section "Input color coding") and, presumably, other small molecules (*52*). We previously showed that addition of BSA does not affect bacterial growth (*33*).

A bank of kChip-deployable carbon source growth substrates was developed from which libraries could be readily created. Carbon compounds in this bank met the following criteria: the compounds were soluble at 2% w/v in water; the solutions were emusifiable using Bio-rad QX200 cartridges; the integrity of the color code dye signals were maintained despite the presence of the carbon compound (see section "Input color coding"). A total of 33 carbon compounds were chosen that passed these criteria, representing a diversity of growth substrates including monosaccharides, oligosaccharides, polysaccharides, carboxylate ions, amino acids, sugar alcohols, and a nucleic acid (**fig S2**).

In the coculture experiment, a total of 40 environmental conditions were used. These included the 33 chosen compounds at 0.5% w/v; five of these compounds (glucose, glycerol, pyruvate, proline, and sucrose) at 0.05% w/v; an even mix of all 33 compounds (totaling 0.5% w/v); and a no-carbon control.

Carbon source plates consisting of the 40 carbon source conditions in water (4X experiment concentration) were frozen at -20°C and thawed at the onset of each round of the experiment (see section "kChip coculture construction"). We previously demonstrated preservation of the frozen carbon substrate plates by showing tight correspondence in the growth of *Escherichia coli* K-12 MG1655 on the freshly prepared carbon substrate plate and plates stored in -20C for 3 days and 15 days (*33*).

Microbial culturing

All labeled and unlabeled monocultures initially underwent two pre-experiment regrowth cycles ("starter phase" in a rich medium and "preculture phase" in minimal medium) and, at the onset of the experiment ("experiment phase"), were normalized to a starting density of $OD_{600}$ = 0.02 in carbon-less minimal medium.

In the first regrowth cycle, the starter phase, glycerol stocks of the unlabeled and labeled strains were inoculated into 525 µL (0.8-mL-deep 96-well plate) of Lysogeny broth (LB) medium (25°C, 220 RPM, 16 hr). Inoculations from glycerol stocks were conducted via pin replicator (sterilized via 70% v/v ethanol bath and heat treatment between inoculations). The second regrowth cycle, the preculture phase, began with washing all cultures in carbon-less M9 medium two times and then diluting cultures (1/50) into 1 mL M9 medium with 0.5% w/v glucose (25°C, 220 RPM, 24 hr). Finally, the experimental phase began by washing cells three times in a carbon-less M9 medium to remove residual glucose and normalizing to a starting $OD_{600}$ of 0.02 (or ~20 cells/droplet depending on the strain). We previously have shown that bacteria grow on kChip with growth dynamics similar to standard culture platforms like Erlenmeyer flasks and microtiter plates (*33*, *34*).

Input color coding

Every unique input to the coculture screen (*e.g.* a bacterial culture or environmental condition) received a "color code", or unique ratio of three fluorescent dyes (standardized to a total final dye concentration of 2.5 μM), prior to generating droplets (**fig S1B**). Each set of three dyes collectively labeled each specific input. The dyes included Alexa Fluor 555, Alexa Fluor 594, and Alexa Fluor 647, all of which have distinct excitation and emission spectra and dyes did not interfere with GFP. We previously showed that the color coding dyes do not affect bacterial growth (*33*).

Droplet preparation and pooling

Each 1-nL droplet contained either a labeled/unlabeled pair or a single carbon source (**Fig 1A**). The labeled strain was projected to each unlabeled strain (1:1 ratio) just before making droplets. Droplets were produced on a Bio-Rad QX200 Droplet Generator (which generated roughly 20,000 ~1-nL emulsifications prepared per 20 μL input for eight inputs at a time; three minutes per 8-input cartridge). The continuous phase was a fluorocarbon oil (3M Novec 7500). For droplet making, 2% w/w fluorosurfactant (RAN Biotech 008 FluoroSurfactant) was added to stabilize droplets.

For each kChip loading (see section "kChip coculture construction"), about 5,000 droplets per each of the labeled/unlabeled inputs (22 + 2 empty controls, or a total of 24 ✕ 5,000 = 120,000 droplets) and about 3,000 droplets per carbon source (39 + 1 empty control, or a total of 40 ✕ 3,000 = 120,000 droplets) were pooled, generating a total of about 240,000 droplets where half contained cultures and the other half contained carbon sources. As a result, about a half of randomly generated pairwise droplet combinations on the kChip contained one labeled/unlabeled pair (premerge $OD_{600}$ = 0.04 for each strain; post-merge $OD_{600}$ = 0.02 for each strain;) and one carbon source droplet (final concentration = 0.5% w/v or 0.05% w/v).

kChip design

All kChips were designed in AutoCAD (Autodesk). Each kChip (62 mm ✕ 72 cm) possessed the following features: (a) An array of ~34,000 microwells (50 mm ✕ 60 mm microfluidic field) where each microwell (slot-shaped, 148.2 μm across, 296.4 μm long) was designed to group $k$ = 2 droplets and arrayed with 50-μm inter-microwell spacing; (b) Internal posts within these microwells designed to (i) reduce overfilling (via droplets squeezing into a microwell), (ii) reduce underfilling (via droplets exiting microwells due to the oil flow associated with the kChip loading procedure), and (iii) inhibit the entry of large droplets inherent to the droplet pool (*i.e.* a low-pass size filter); (c) A series of 30 90-μm deep moat-like slots designed to trap small droplets (*i.e.* a high-pass size filter) spaced 50 μm apart from each other, 400 μm from the onset of the microfluidic field, and 3 mm inset from the edge of the kChip; and (d) A loading slot into which droplets were injected via micropipette.

The optimal microwell geometries and distances between posts was optimized based on the choice of medium and concentration of fluorosurfactant (RAN Biotech 008 FluoroSurfactant), which we have previously shown affect the size of droplets produced by a Bio-Rad QX200 Droplet Generator, and by extension, droplet grouping and merging performance (*33*).

Photomasks were generated from AutoCAD designs (FineLine Imaging). kChip designs were then fabricated to 110-120 μm feature height using photolithography on silicon wafers

(Microchem SU8-2050). Microwells produced from this feature height were found to best trap droplets in a monolayer, as deeper features can allow droplets to stack causing loading of an undesired number of droplets (*33*). These wafers were then embedded into custom molds to create PDMS (Dow Corning Sylgard) kChips by soft lithography with consistent thickness (0.635 cm) and droplet-loading slot location and size. The side of the kChip that contained microwell features was then coated with 1.5 μm parylene C by vapor deposition (Paratronix) to inhibit water loss from droplets and stiffen the kChip to prevent interior collapse during droplet loading.

<u>kChip coculture construction</u>

The following steps were performed for each kChip, with 6 kChips run in sequence per day (for a total of 24 kChip run over 4 days, with 2 kChips repeated due to poor droplet loading). As previously described, a single labeled strain was projected to all unlabeled strains, each labeled/unlabeled input and each carbon source input received a unique color code, droplets were made from all inputs, and all droplets were pooled together (**Fig 1A**, Steps 1-2). With the assistance of a loading apparatus (**fig S2A**), the droplets were loaded onto a kChip in one pipetting step (**Fig 1A**, Step 3).

A kChip loading apparatus, which was required to load droplets into microwells, consisted of a "loader top" and "loader base" (**fig S1A**). The loader base held in place a piece of custom-cut glass (Brain Research Laboratories; 1.2 mm thickness) made hydrophobic via pretreatment with Aquapel. The top side of the kChip, which was not coated with parylene, spontaneously formed a seal with the loader top. Four neodymium magnet pairs were oriented such that the two acrylic pieces repelled each other. Working against this repulsive force, the loader top was lowered toward the loader base via tightening nuts until the desired standoff between the glass and kChip was attained (~500-700 μm) to create a space for flow under the microwells. Via an open slot passing through the loader top and kChip, the flow space was pre-wetted with an injection of oil (~3 mL to fill the entirety of the flow space) followed by the pooled droplets. Each kChip consisted of an array of microwells, each designed to randomly group a specific number of droplets. Here we used only "$k = 2$" microwells that each group two droplets. Buoyant in the surrounding oil, the droplets were distributed around the flow space via tilting the loading apparatus. After the droplets had passed through the flow space and entered the $k = 2$ microwells, additional oil (no fluorosurfactant) was flushed through the device to wash away excess droplets and fluorosurfactant. The kChip, coupled to the top piece, was removed, and sealed with a transparent PCR film (*35*), limiting inter-microwell crosstalk (*34*). The edges of the film were trimmed off. A second set of four neodymium magnets were used to couple to the top piece/kChip to a microscope stage adapter.

Setting up the kChip loading apparatus in preparation to receive droplets took 5-10 minutes and was completed ahead of time. The remaining setup time was ~30 minutes: Droplet making took ~3 minutes per eight inputs on the Bio-Rad QX200, droplet pooling and mixing took ~5 minutes, loading the kChip took ~5-10 minutes, and scanning the kChip took ~12-15 minutes.

The kChip was scanned initially at 2X magnification to identify the droplets in each microwell from their color codes (**Fig S1B**). Droplets were then merged within their microwells via exposure to an alternating current (AC) electric field (4.5 MHz, 10,000-45,000 volts) underneath the PCR film (**Fig 1A**, Step 4, **Fig S1B**). The field was generated by a corona treater (Electro-Technic Products), the tip of which was moved around the PCR film for ~10 seconds. Without application of the electric field or undue physical force, spontaneous merging of droplets was rare (detected as incorrectly loaded microwells in **Fig S1B**). The kChip was imaged subsequently to measure the yield of the labeled strain, defined as the GFP signal at 0, 24, and 72 hr (**Fig 1A**, Step 5, **fig S1B-C**).

Of the 22 kChips that were loaded well, two were removed downstream at the data analysis stage due to low GFP signal of the labeled strain (and cocultures containing these strains' unlabeled versions across the other kChips were also removed from the data analysis). We previously showed there is minimal evaporation in droplets between 24 and 72 hrs (*33*), and expect this has little effect on microbe growth given our observation that different microbes reach saturation in droplets and 96-well plate cultures on similar timescales.

We developed an image analysis pipeline, previously described (*33*) to: (a) Identify droplets as circular objects within the image; (b) Decode the contents of each droplet based on the color code; (c) Assign each droplet to a microwell; (d) filter out poorly loaded microwells from the data; and (e) Measure the average fluorescence of the merged droplets in each microwell.

Across the 20 kChips included in the data analysis, a total of 16,000 unique labeled/unlabeled/environment combinations were possible ([20 monocultures + (190 $\times$ 2) cocultures] $\times$ 40 environments = 16,000 combinations) (**Fig 1B**). After quality filtering data through our image analysis pipeline, the dataset contained 180,408 data points, with each of these combinations appearing 10.3 times on average and 97% appearing $\geq$3 times (**fig S4**). By integrating data among kChips, bidirectional pairwise interactions were deduced (**Fig 1C, fig S5, S6**).

Fluorescence imaging

All fluorescence microscopy was performed using a Nikon Ti-E inverted fluorescence microscope with fluorescence excitation by a Lumencor Sola light emitting diode illuminator (100% power setting). Images were taken across four fluorescence channels—three for the color codes and one additional channel for fluorescence-based assays. Each dye and the assay signal was detected with a different excitation wavelength generated by a collection of excitation filters: GFP by Semrock GFP-1828A (blue excitation); Alexa Fluor 555 dye by Semrock SpGold-B (green excitation); Alexa Fluor 594 dye by Semrock FF03-575/25-25 [excitation filter] + FF01-615/24-25 [emission filter] (yellow excitation); and Alexa Fluor 647 dye by Semrock LF635-B (red excitation). The emission signals corresponding to each dye channel were used to identify the contents of a given droplet within each droplet grouping prior to droplet merging (**Fig 3B)**. The final channel was used post-merge and at subsequent time points to quantify the assay signal (**Fig 3B)**.

Bootstrap resampling and interaction classification

To measure $E_{B\rightarrow A}$, the effect of unlabeled strain B on labeled strain A for a given carbon source environment, the $\log_2$ of the ratio of its yield of A in coculture (median of $A_{+B}$ replicates) to monoculture (median of $A_{mono}$ replicates) was calculated (**Fig 1D**, **fig S5A**). In instances where either of these values fell below the detection limit (DL), they were replaced with DL, which was calculated as the 90% percentile of the distribution of $A_{+B}$ when no carbon source was present. Several examples of this calculation are provided in **fig S5B**. Positive values indicated facilitation, negative values indicated inhibition, and 0 indicated no detected effect.

The coordinate ($E_{B\rightarrow A}$, $E_{A\rightarrow B}$) represented both sides of an interaction. To qualitatively classify this interaction, this point was plotted on a Cartesian plane (and, as necessary, reflected to the left of the identity line $y = x$). Uncertainty was calculated via bootstrapping: 1000 calculations were performed for $E_{B\rightarrow A}$ and $E_{A\rightarrow B}$ via resampling $A_{+B}$, $A_{mono}$, $B_{+A}$, and $B_{mono}$. The 25th and 75th percentiles of the resulting distributions were plotted (**Fig 1E**, **fig S6A**). An interaction was classified as mutualism (+,+) if both sets of uncertainty bars fell within the first quadrant; as a parasitism if they both fell within the second quadrant (+,–); and as a competition if they both fell within the third quadrant (–,–). Error bars passing over quadrants indicated that a classification for at least one of the two effects did not adequately separate from no effect. If an uncertainty bar passed over the *y*-axis, an interaction was classified as a commensalism (+,0); if an uncertainty bar passed over the *x*-axis, the interaction was classified as an amensalism (–,0); if both uncertainty bars passed over the *x*- and *y*-axes, the interaction was classified as a neutralism (0,0). Examples of these pairwise classifications are shown in **fig S6A**.

The point ($E_{B\to A}$, $E_{A\to B}$) occupied a radial continuum of possible pairwise interactions, with the magnitude m and the angle Θ of this point in polar coordinates providing a quantitative description of the interaction strength and type, respectively (**Fig 1F**). The value Θ specifically represented the relative size of the effects of two strains on each other. To determine Θ, the line $y = -x$ was assigned to 0° (representing a balanced parasitism of equal and opposite effects). Values $-90° < \Theta < -45°$ quantified competition, with –90° indicating that two strains inhibited each other equally; –45° indicated amensalism; $-45° < \Theta < 0°$ quantified parasitism where the inhibitory effect outweighed the facilitative effect; $0° < \Theta < 45°$ quantified parasitism where the facilitative effect outweighed the inhibitory effect; 45° indicated commensalism; and $45° < \Theta < 90°$ quantified mutualism, with 90° indicating that two strains facilitated each other equally. The distribution of m and Θ are given in **fig S8A** (and separated by carbon source in **fig S8D**). Θ for all pairwise interactions is hierarchically clustered in **fig S9**.

For a given set of interactions (*e.g.* all pairs among a phylogenetic group) the mean interaction [$\mu(E_{B\to A})$, $\mu(E_{A\to B})$] was sometimes calculated (**fig 6SB**). The average interaction magnitude m and average interaction type Θ of this average interaction were also calculated. The variance in a set of interactions can also be calculated as an interaction diversity metric. This analytical framework was applied to the entire dataset (**Fig 2**, **fig S7**) and to subsets of the data organized by properties of the environments, such as biochemical classification (**Fig 3B**), or properties of strains, such or phylogenetic distances (**Fig 3D**). The distribution of the average m and average Θ for interactions grouped by coculture pair and by carbon source are given in **fig S8B** and **fig S8C**, respectively.

<u>Binning interactions by metabolic distance or monoculture growth</u>

Using measured resource utilization profiles (monoculture growth values across all carbon sources normalized to the maximum growth per strain, per time point) (**fig S2**), the Euclidean distance between each resource utilization profile was computed for each strain pair (**fig S3C**) as a measure of metabolic similarity. Whereas the phylogenetic distances among pairs produced a bimodal distribution that reflected the two taxonomic orders, the metabolic distance values produced a continuous bell-shaped distribution (**fig S10**).

We binned the 190 possible cocultures into 8 discrete metabolic distance bins of increasing dissimilarity and counted interaction types in each bin across all carbon sources (**Fig 3E**, **fig S11**). (Cocultures of a labeled strains cocultured with their unlabeled counterparts, for which the metabolic distance was 0, were grouped as Bin #0.) In the center of the metabolic distance distribution (Bins #3, #4, #5, and #6), each bin spanned 0.25 Euclidean distance units. Because there were fewer data points nearer the tails of the distribution, Bins #2 and #7 each spanned 0.5 Euclidean distance units, and Bins #1 and #8 spanned 1 Euclidean distance unit. The resulting bins each had roughly equal numbers of data points.

Unlike binning by metabolic distance, binning by monoculture growth disregarded any larger interaction patterns within a given strain pair, i.e. each interaction generated among a pair across each carbon source was independently binned only by the degree to which each strain grew on the given carbon source. Normalized monoculture yields (**fig S2**) were placed into one of seven bins (cutoffs: 0, 0.005, 0.05, 0.1, 0.2, 0.3, 0.5, 1). The first bin, [0 0.005), represented undetectable growth (within background noise) as qualified as “no growth” in analyses of obligate facilitation (**Fig 4D**). With the exception of the second bin, [0.005 0.05), which spanned a decade, bins cutoffs were roughly based on exponential doublings.

<u>Interaction networks and binning</u>

Interactions networks were constructed for all pairwise interactions occurring per carbon source (examples in **Fig 4A** and all networks in **Fig S13**). The nodes, each representing a strain, were arranged concentrically by carbon utilization similarity (same order as in **fig S2**). The size of the node corresponded linearly to the normalized monoculture growth (**fig S2**). Edges between nodes, each representing a pairwise interaction, were colored by interaction classification (with neutralisms not shown). The thickness of the edge corresponded to interaction strength m.

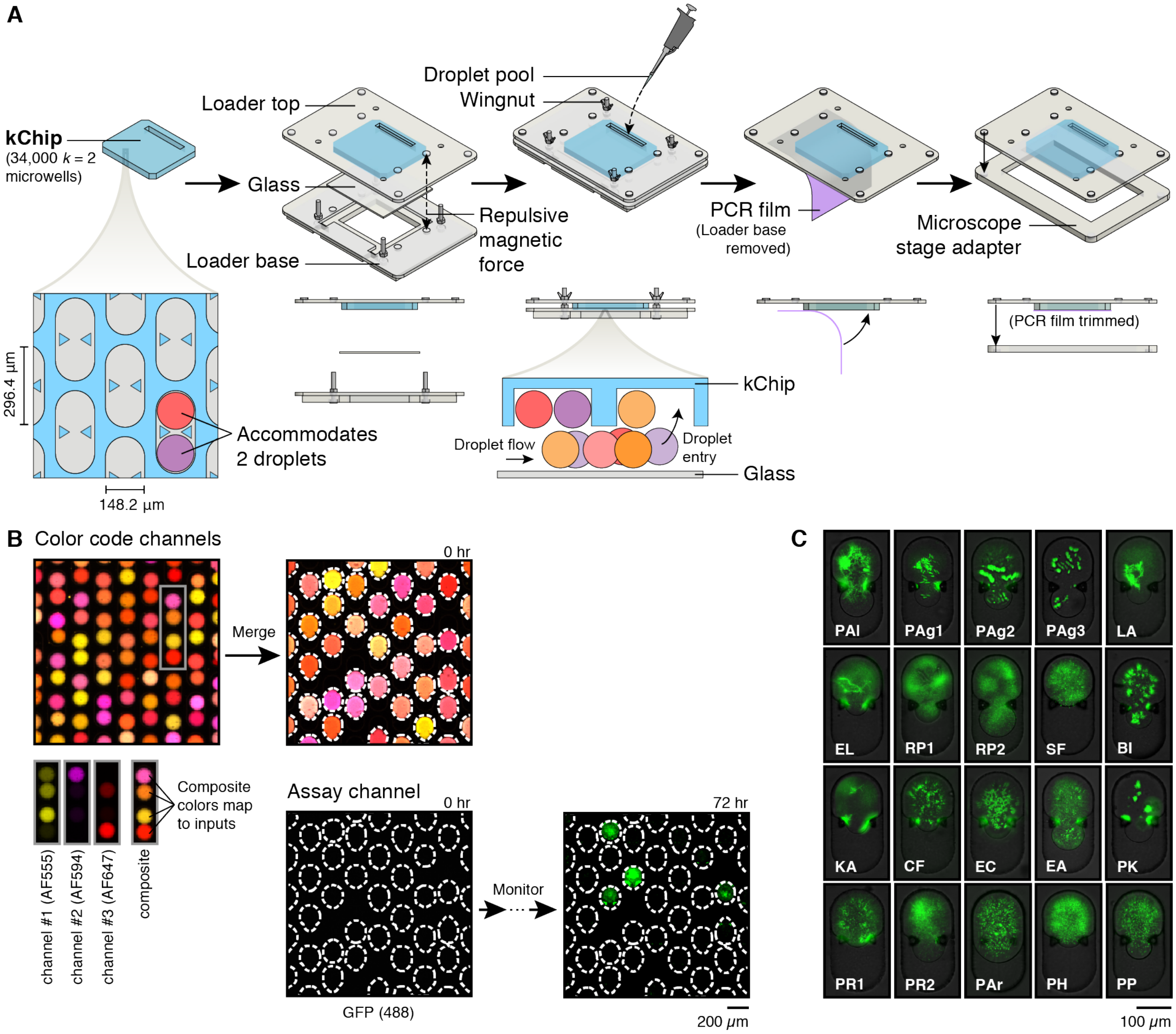


**Fig S1. kChip experimental workflow. A.** A kChip is chosen with a desirable microwell array layout, e.g. all $k = 2$ microwell geometries designed to capture two droplets each. Positioned inside of an acrylic loading apparatus, the kChip is suspended via repulsive magnetic interactions over a glass substrate to create a droplet flow space. Droplets are pipetted into a loading slot and enter microwells as they move through. After loading droplets, the kChip is sealed with a PCR film. **B.** The kChip is initially imaged in multiple channels (2X magnification) to determine the identity of each droplet. Here a specific ratio of three Alexa Fluor fluorescent dyes (AF555, AF594, and AF647) is used to barcode each droplet; the composite color indicates the droplet identity. After merging the droplets, an additional assay channel is used, e.g. GFP, to measure growth over time. **C.** Images of labeled strains in example cocultures (with arbitrary unlabeled strain and carbon source) (10X magnification). Full strain names are provided in **Table S1**.

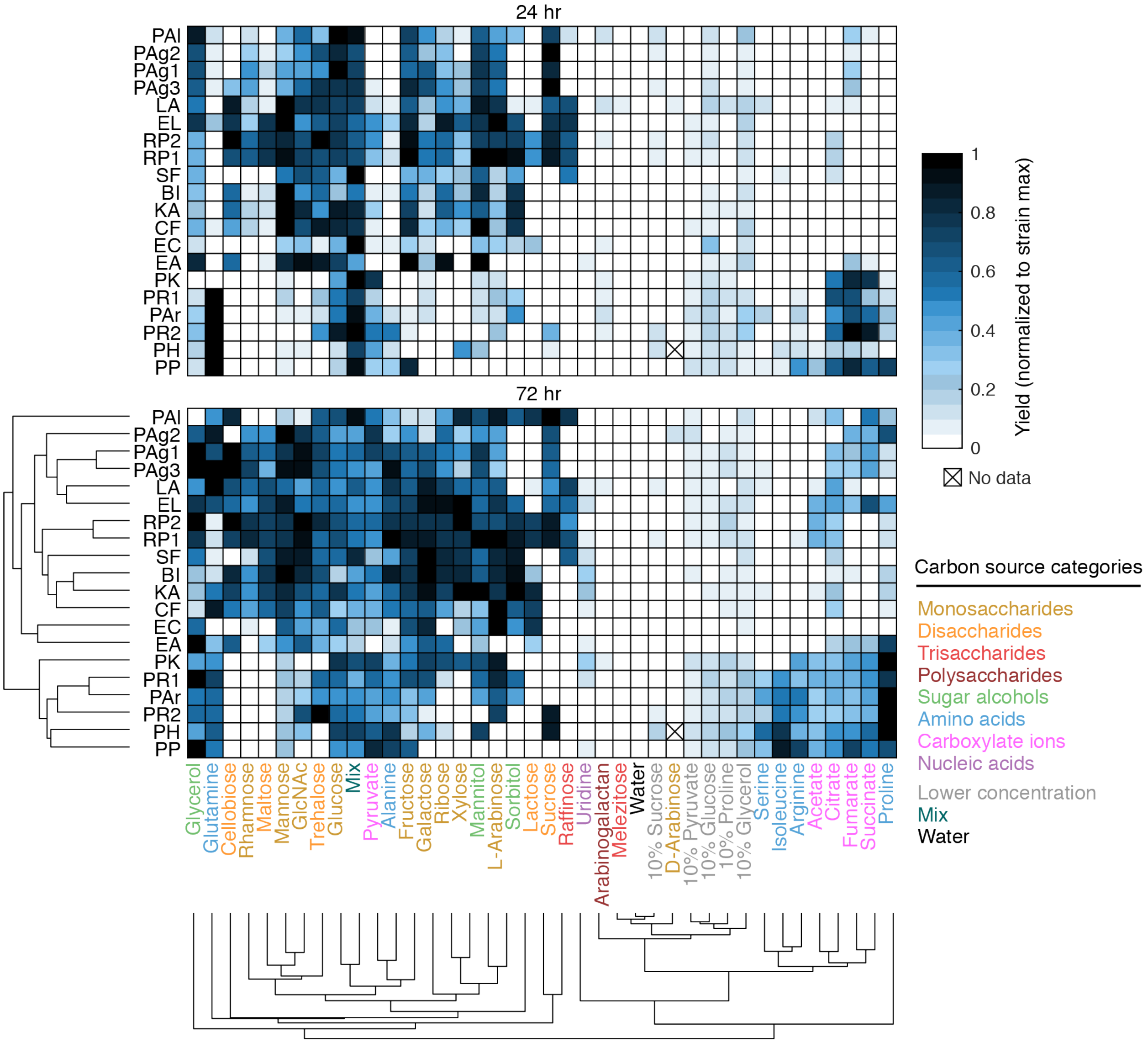


**Fig S2. Carbon source utilization profiles of soil bacterial strains.** Each strain's ability to grow on each carbon source was determined from kChip microwells containing a labeled strain in monoculture and a carbon source. Yield values (median GFP measurement across replicates) were background-subtracted (background = median yield of labeled strain with no carbon) and normalized to the maximum yield value observed (per strain per time point). Carbon sources and strains were hierarchically clustered at the 72 hr time point.

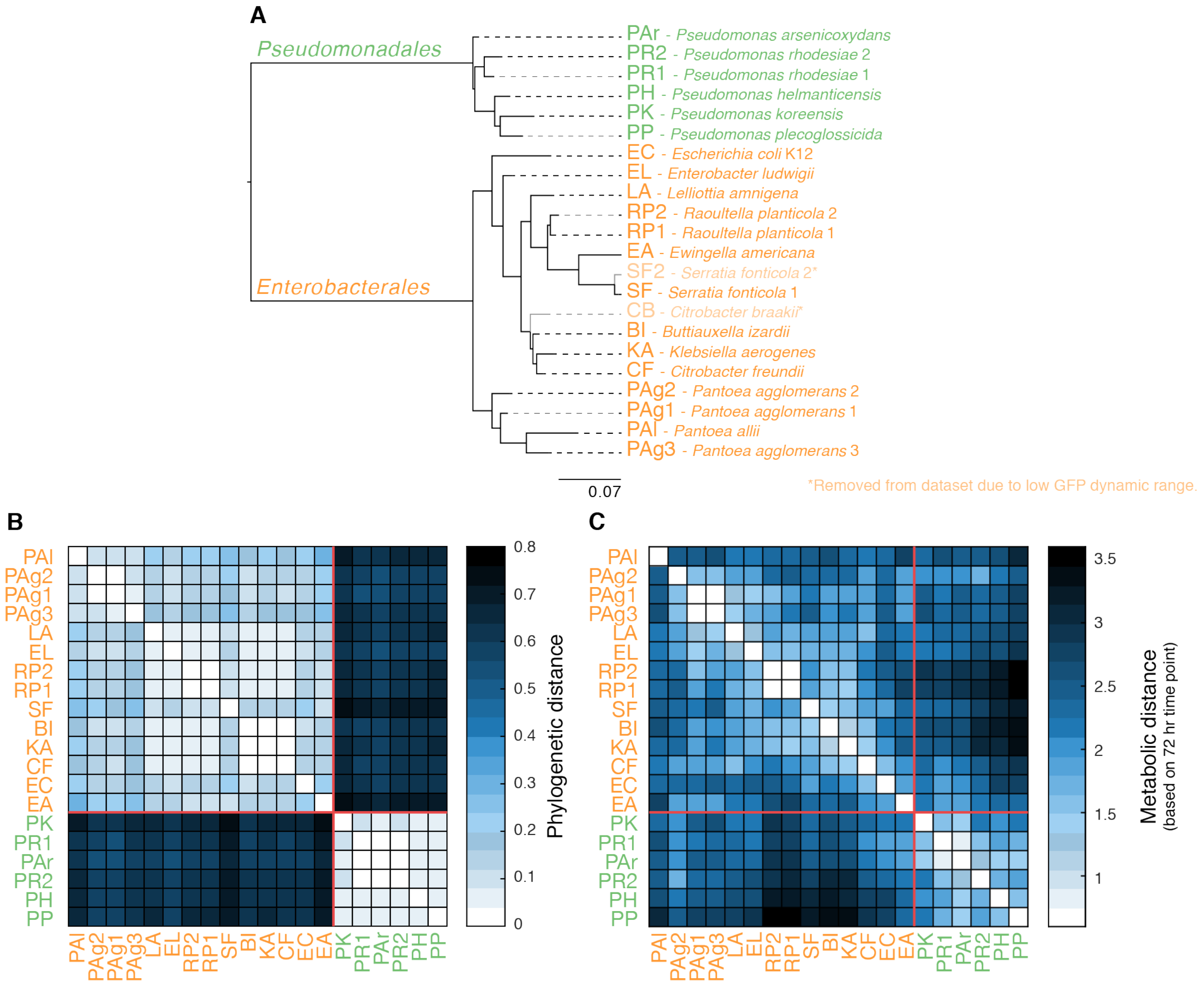


**Fig S3. Strain pairwise comparison metrics. A.** Phylogenetic tree built with maximum likelihood estimate, utilizing alignment of full 16S sequences. Sequences of the full 16S rRNA gene (~1500bp) were obtained via sanger sequencing. Sequences were trimmed to quality threshold ≥ 20 before merging the forward and reverse reads of each bacterial 16S. The sequence for *E. coli* K12 was obtained directly from NCBI. *Sulfolobus solfataricus*, a thermophilic archaeon, was used in the phylogenetic reconstruction as an outgroup species to root the tree. MUSCLE with default parameters was used to align the sequences. PhyML-SMS with default parameters was used to select GTR+G+I as the best model and to infer the tree. Units are 'Substitutions per base pair of the 16S gene. **B.** Pairwise phylogenetic distances between strains were calculated directly from the phylogenetic tree with R (Library Phytools, command 'drop.tip'). **C.** Metabolic distances ordered by carbon source utilization similarity. These distances were calculated as the Euclidean distance between two carbon source utilization profiles (**fig S2**). Red line = separation of two taxonomic families. Orange font = strains of the order *Enterobacterales*. Green font = strains of the order *Pseudomonadales*.

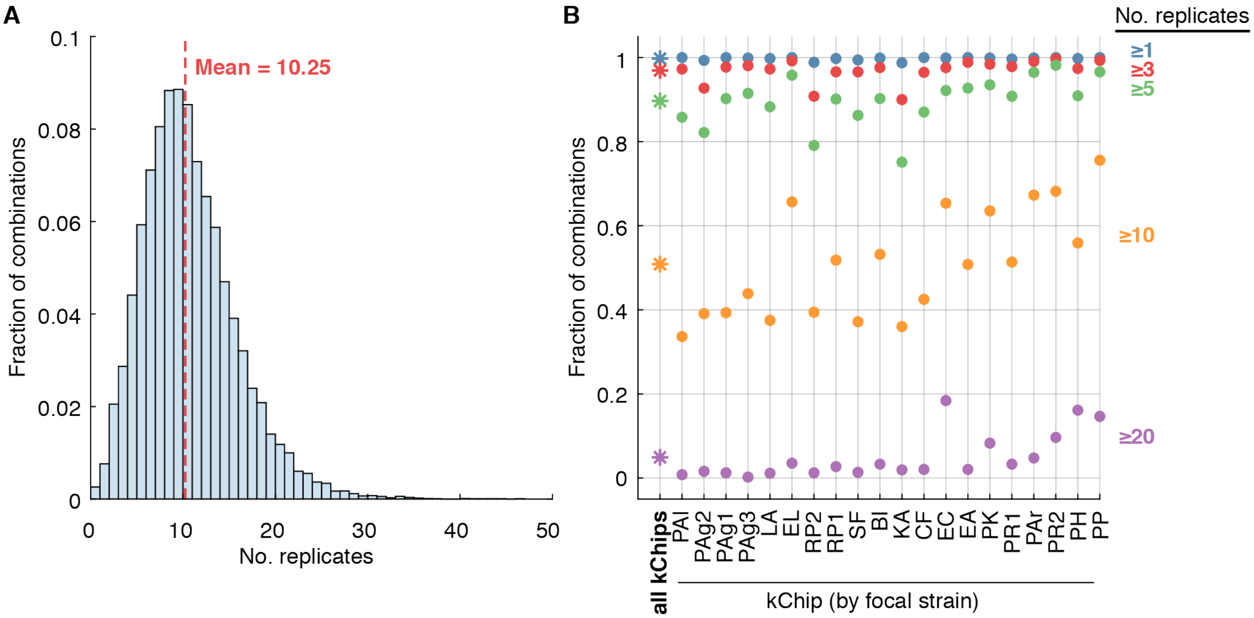


**Fig S4. Combination sampling. A.** Distribution of the number of replicates for all possible combinations of labeled strain/unlabeled strain/environment, including all 180,408 data points and representing all 17,600 possible combinations (i.e. all combinations of 20 labeled strains, [20 + 2 control] unlabeled strains, and [39 + 1 control] carbon sources). **B.** Fraction of combinations represented at least 1, 3, 5, 10, and 20 times for the entire dataset and per kChip.

**A**

$$E_{B\rightarrow A} = \text{Effect of unlabeled B on focal A} = \log_2 \frac{\max(A_{+B}, DL)}{\max(A_{mono}, DL)}$$

$A_{+B}$ = background-subtracted median yield of focal A in coculture with B with given carbon source

$A_{mono}$ = background-subtracted median yield of focal A in monoculture with given carbon source

DL = background-subtracted 90th percentile of A in coculture with B with no carbon source

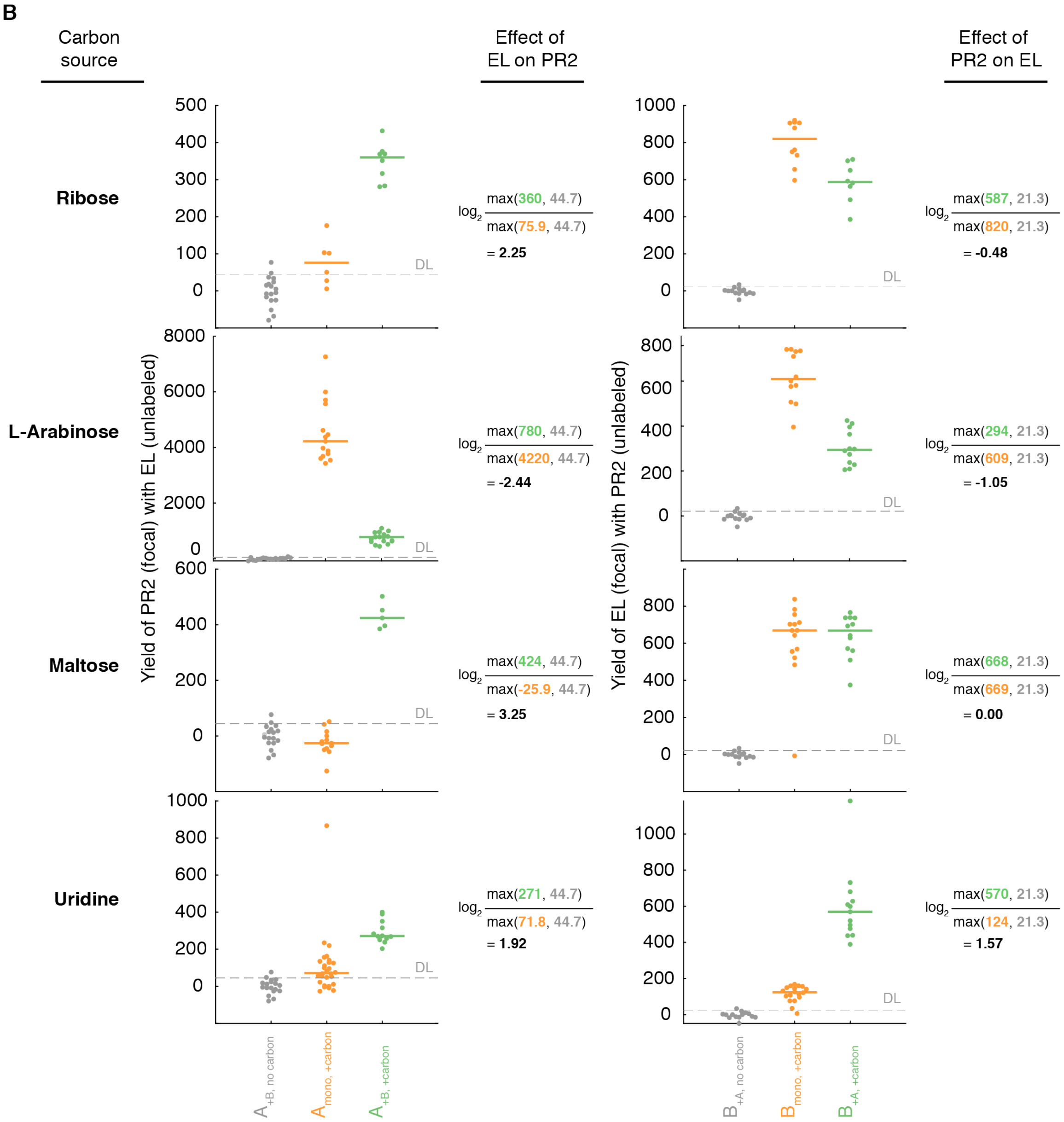


**Fig S5. Interaction effect calculation. A.** Formula for calculating effect of unlabeled strain B on yield of labeled strain A. **B.** Effect calculations for example coculture [*Enterobacter ludwigii* (EL) and *Pseudomonas rhodesiae* #2 (PR2)] on four example carbon sources.

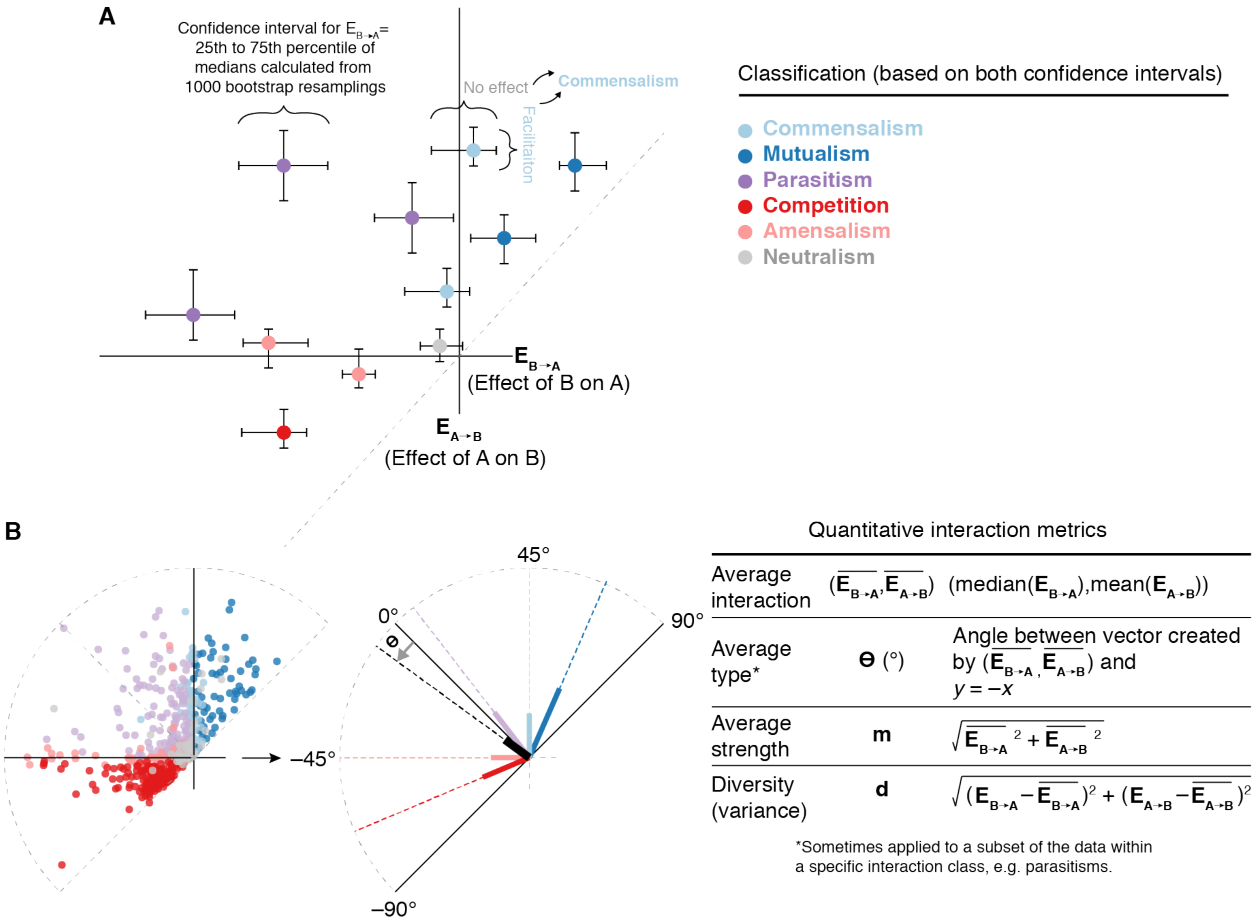


**Fig S6. Pairwise interaction classification and quantitative metrics. A.** Method for assigning pairwise interaction classification (commensalism, mutualism, parasitism, competition, amensalism, or neutralism). Confidence intervals for all one-way effects were generated by bootstrap resampling the effect calculation (**fig S5**). One-way interactions (facilitation, inhibition or no effect) were classified by the effect size and the fraction of the confidence interval that fell within the effect size's quadrant. Two-way interactions were classified by combining two one-way classifications. **B.** Quantitative metrics used to describe pairwise interactions. These include metrics for type, strength, and diversity (with diversity only applying to a set of interactions). The type metric was also used to subset of interactions belonging to specific interaction classifications.

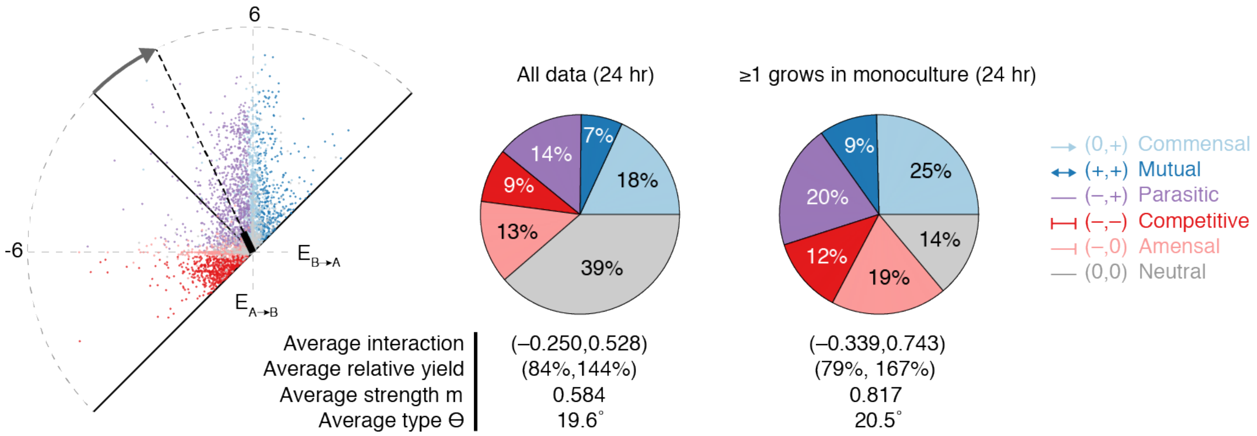


| | All data (24 hr) | ≥1 grows in monoculture (24 hr) |
|---|---|---|
| Average interaction | (–0.250,0.528) | (–0.339,0.743) |
| Average relative yield | (84%,144%) | (79%, 167%) |
| Average strength m | 0.584 | 0.817 |
| Average type ϴ | 19.6˚ | 20.5˚ |

**Fig S7. Interaction distributions for full dataset at 24 hr.** (Left) All pairwise interactions at 24 hr on 33 distinct carbon sources. (Middle) Interaction classification of all data. (Right) Interaction classification excluding cases in which both strains comprising a coculture showed no detectable growth as monocultures on a given carbon source.

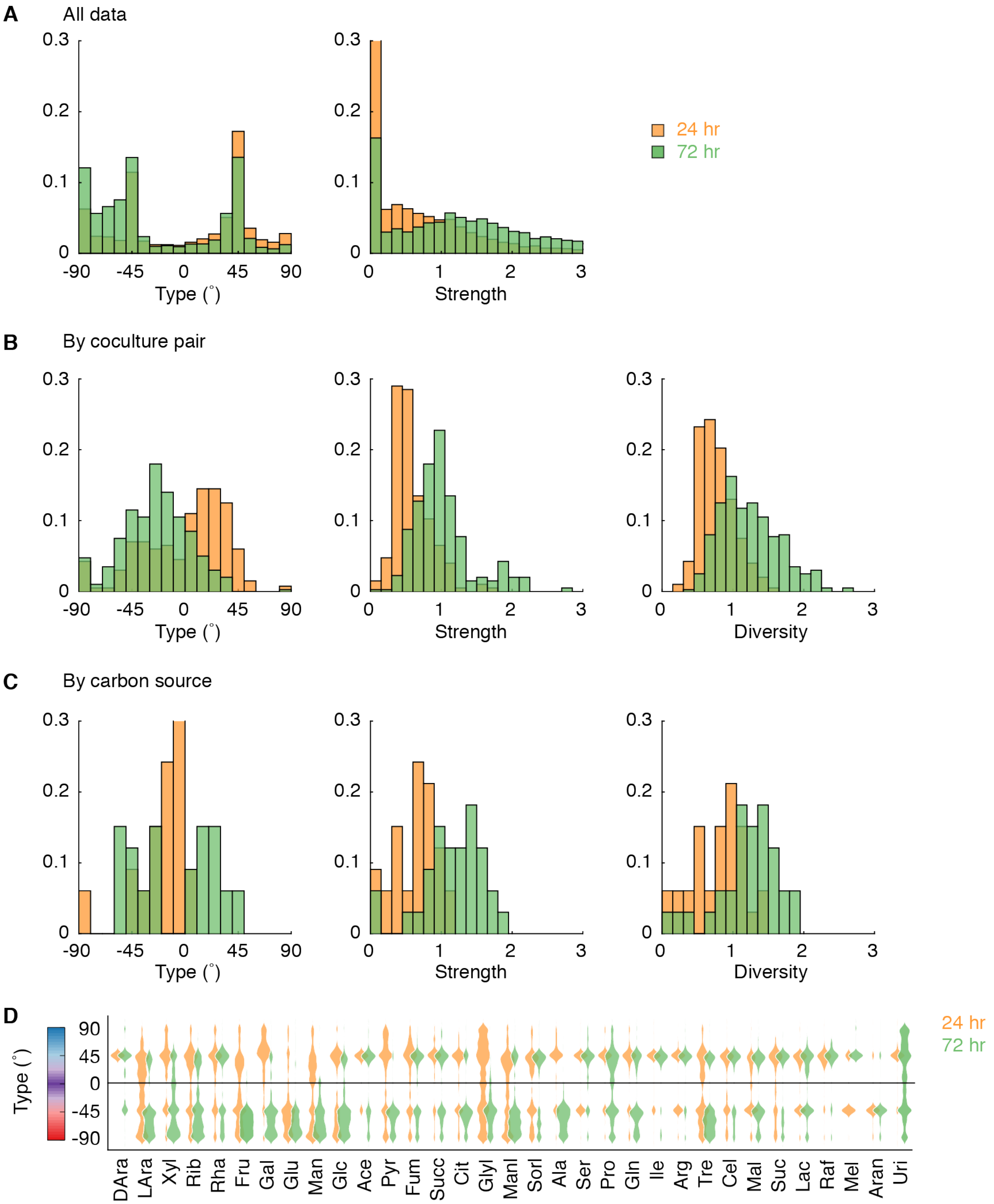


**Fig S8. Interaction distributions by strain pair and by carbon source. A-C.** Distributions of quantitative interaction metrics for all data (A), data averaged by strain pair (B), and data averaged by carbon source (C). Calculations for averages are described in **fig S6. D.** Distributions of interaction type metric organized by carbon source.

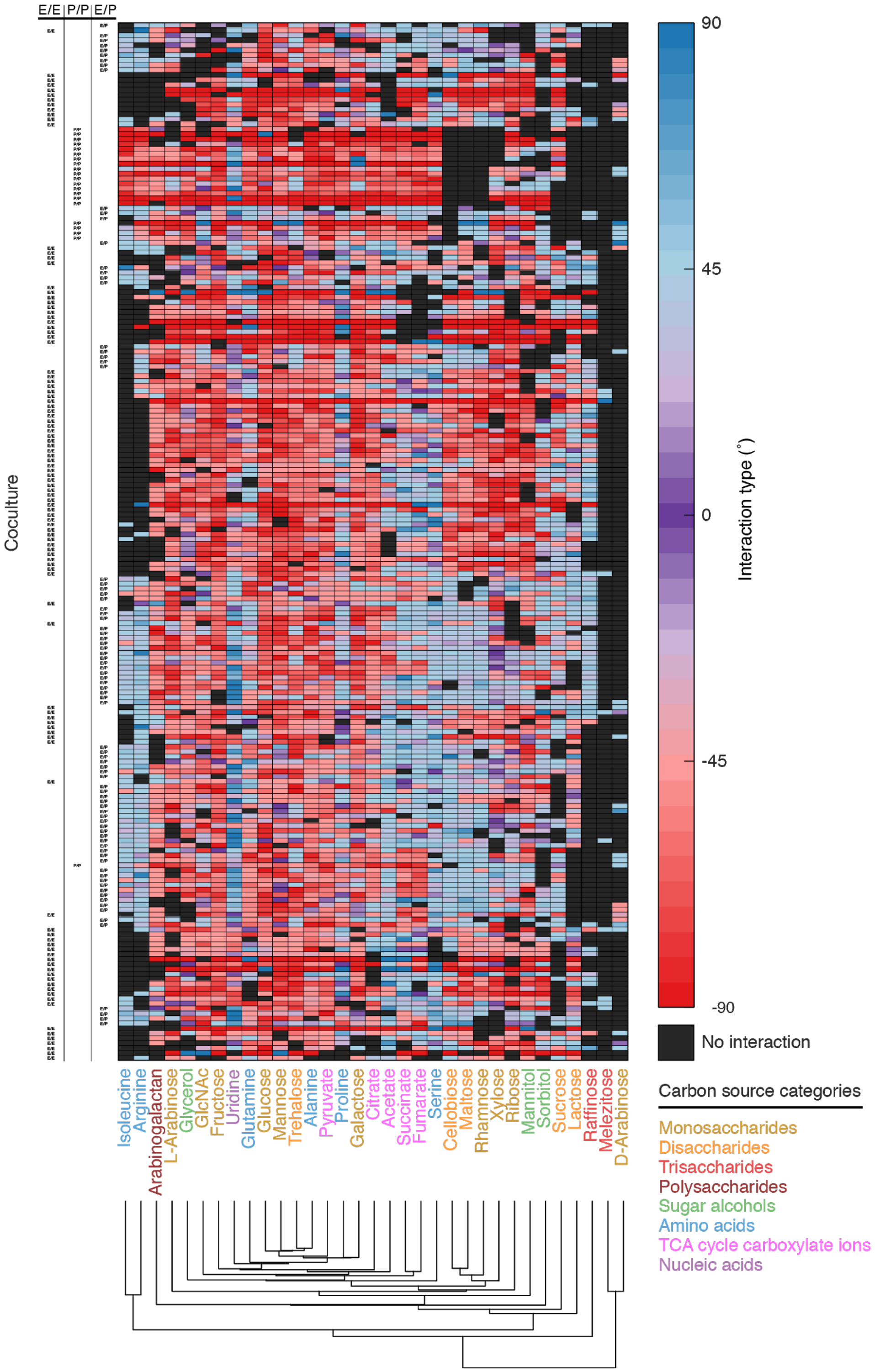


**Fig S9. Interaction types for full dataset.** Strain pairs and carbon sources were hierarchically clustered by interaction type. E = *Enterobacterales*. P = *Pseudomonadales*. All data at 72 hr.

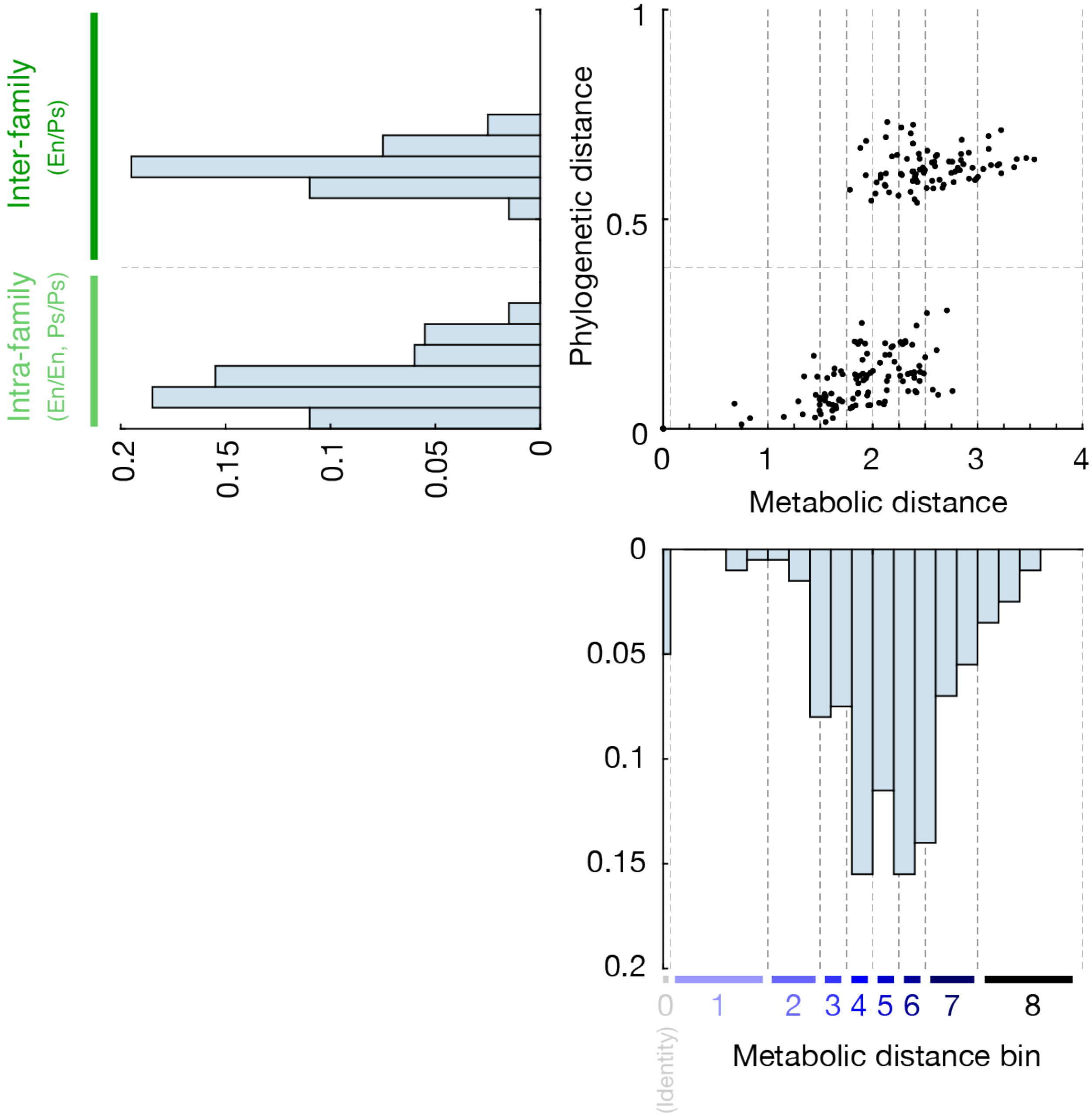


**Fig S10. Binning of phylogenetic and metabolic distances.** Phylogenetic distances occupied a bimodal distribution representing intra-family and inter-family pairs. Metabolic distances occupied a continuous distribution. Metabolic distances bins were drawn to capture roughly even numbers of pairs. In the dense center of the distribution, bins were evenly spaced by 0.25 metabolic distance units (Bins #3-6). The next outermost bins occupied 0.5 metabolic distance units (Bins #2 and #7), and the most outermost bins occupied 1 metabolic distance unit (Bins #1 and #8). Bin #0 represents with-self interactions (with a metabolic distance of 0). En = *Enterobacteriaceae*. Ps = *Pseudomonadaceae*.

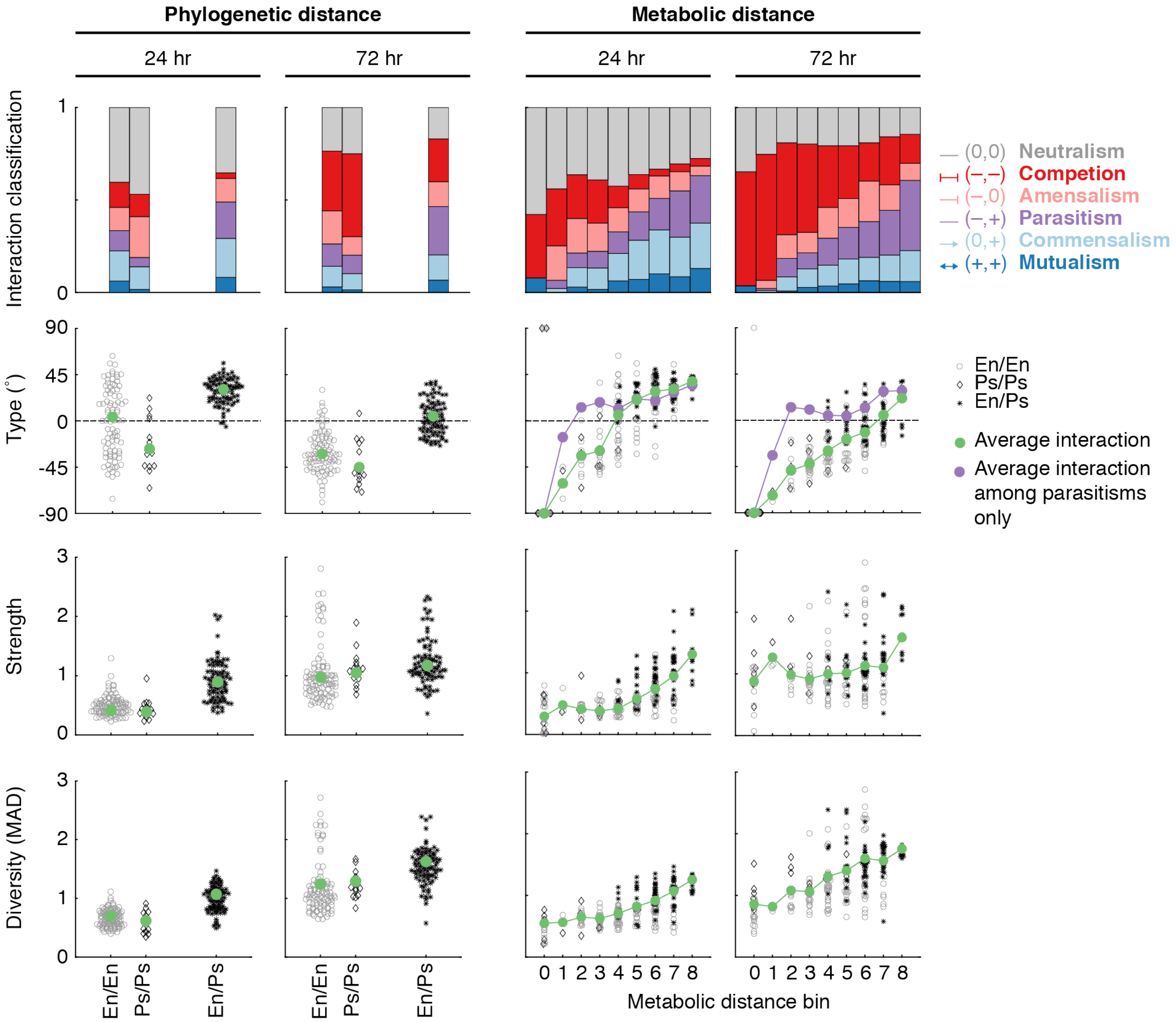


**Fig S11. Interaction metrics by phylogenetic and metabolic distance bins.** Phylogenetic bins represent the intra-family interactions (En/En and Ps/Ps) and inter-family interaction (En/Ps). Metabolic bins represent increasingly metabolically different pairs and are described in **fig S10**.

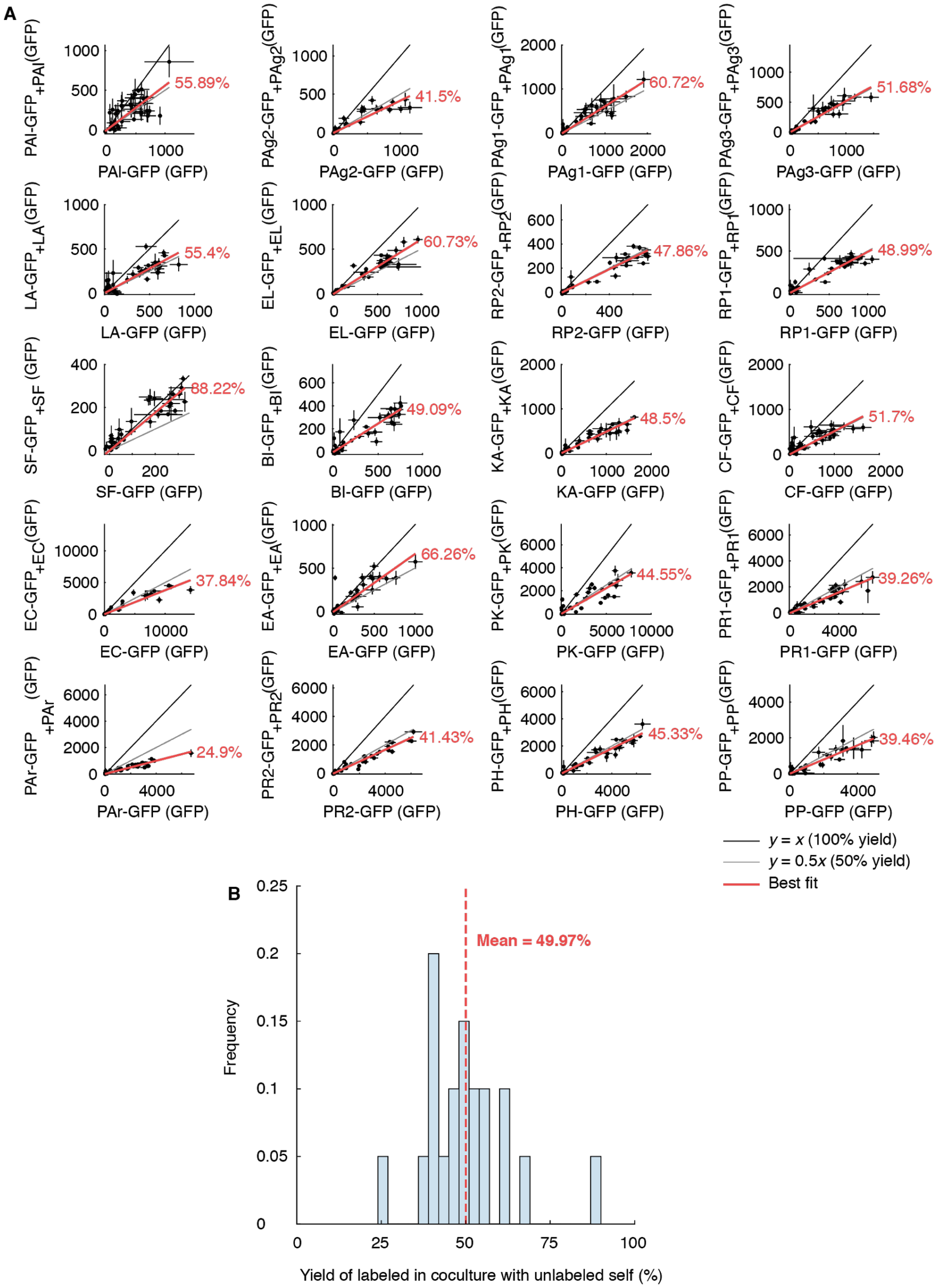


**Fig S12. With-self interactions. A.** Each plot represents an interaction between a labeled strain and its unlabeled counterpart on a given carbon source. The red line indicates the average fraction of the labeled strain relative to its monoculture yield. **B.** Distribution of each strain's average growth with their respective unlabeled self-strains.

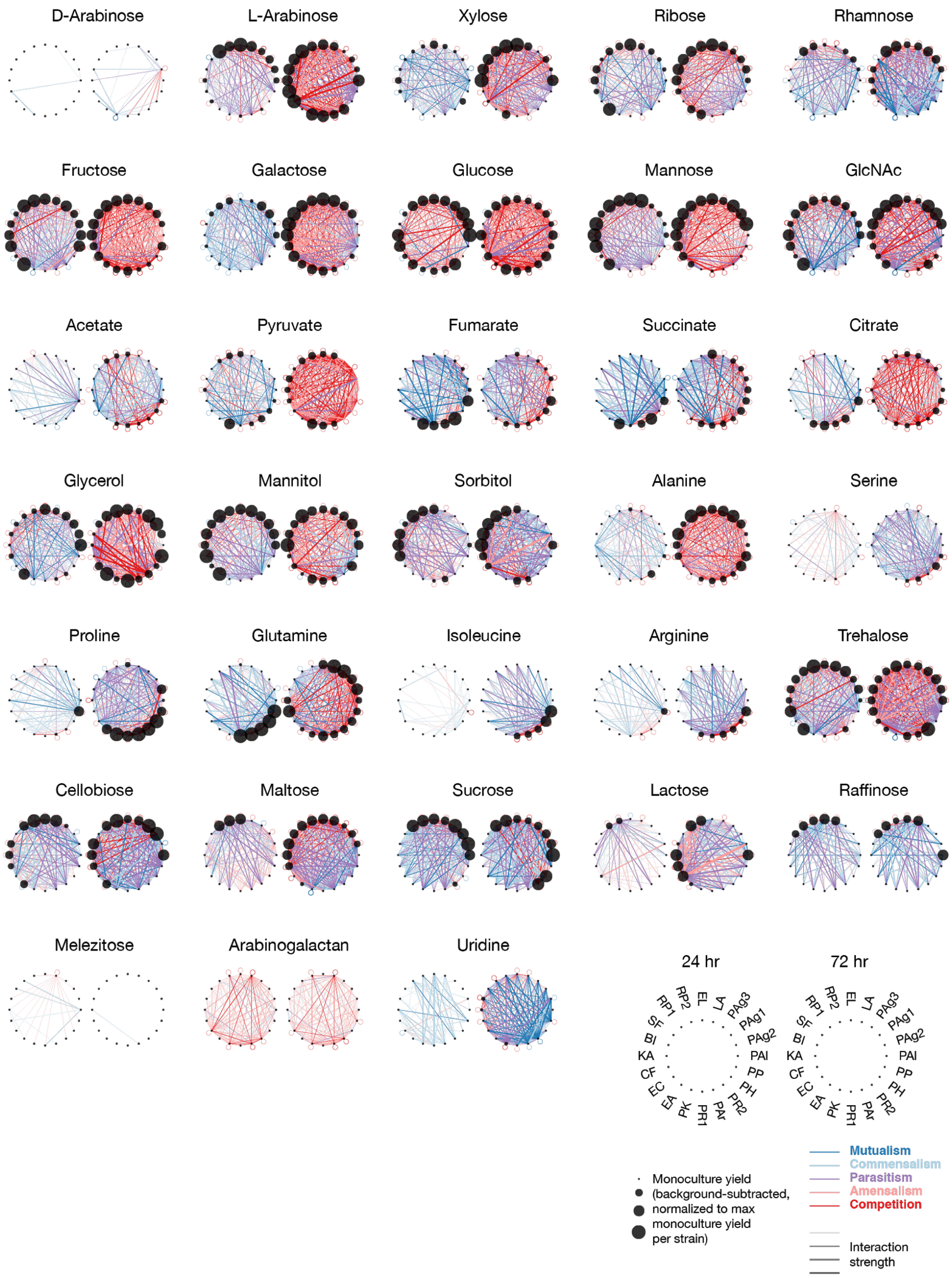


**Fig S13. Interaction networks for each carbon source**. For each carbon source, the pairwise interaction network is plotted at 24 hr (left) and 72 hr (right). The legend (bottom-right) indicates the strain identity of each node. Node size represents monoculture yield (background-subtracted and normalized to maximum monoculture yield per strain). Edge color represents interaction classification. Edge thickness represents interaction strength.

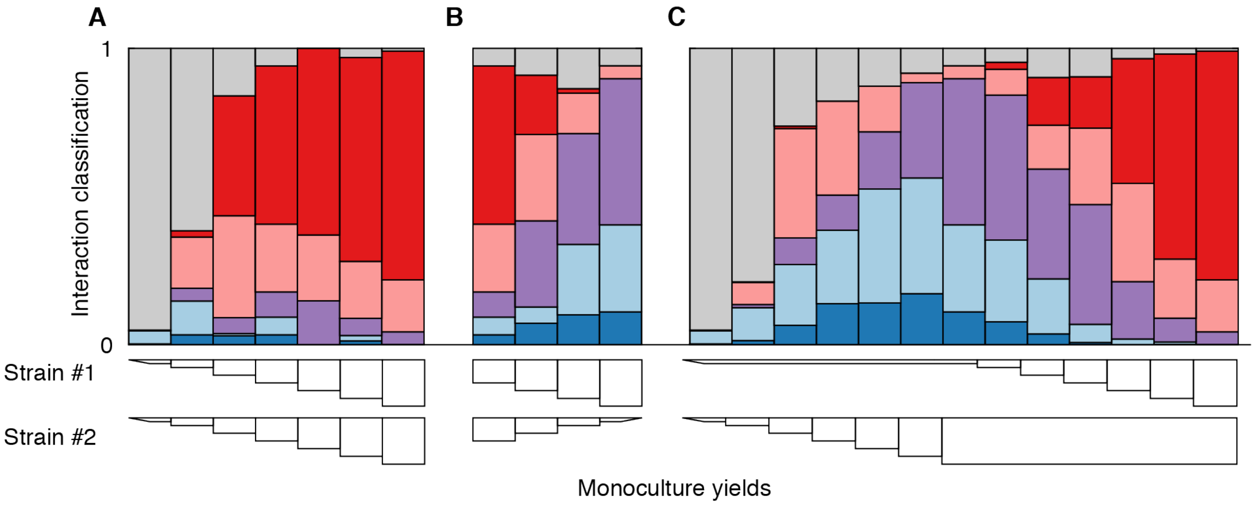


**Fig S14. Interaction classification by monoculture yield comparisons.** Interaction classification when both strains are increasing in monoculture yield across carbon sources (**A**), one strain is increasing and the other is decreasing in monoculture yield (**B**), and one held constant at no-growth or maximum growth while the other strain is increasing in monoculture yield (**C**). Colors indicate interaction classification (with color legend in **Fig 2**). All data at 72 hr.

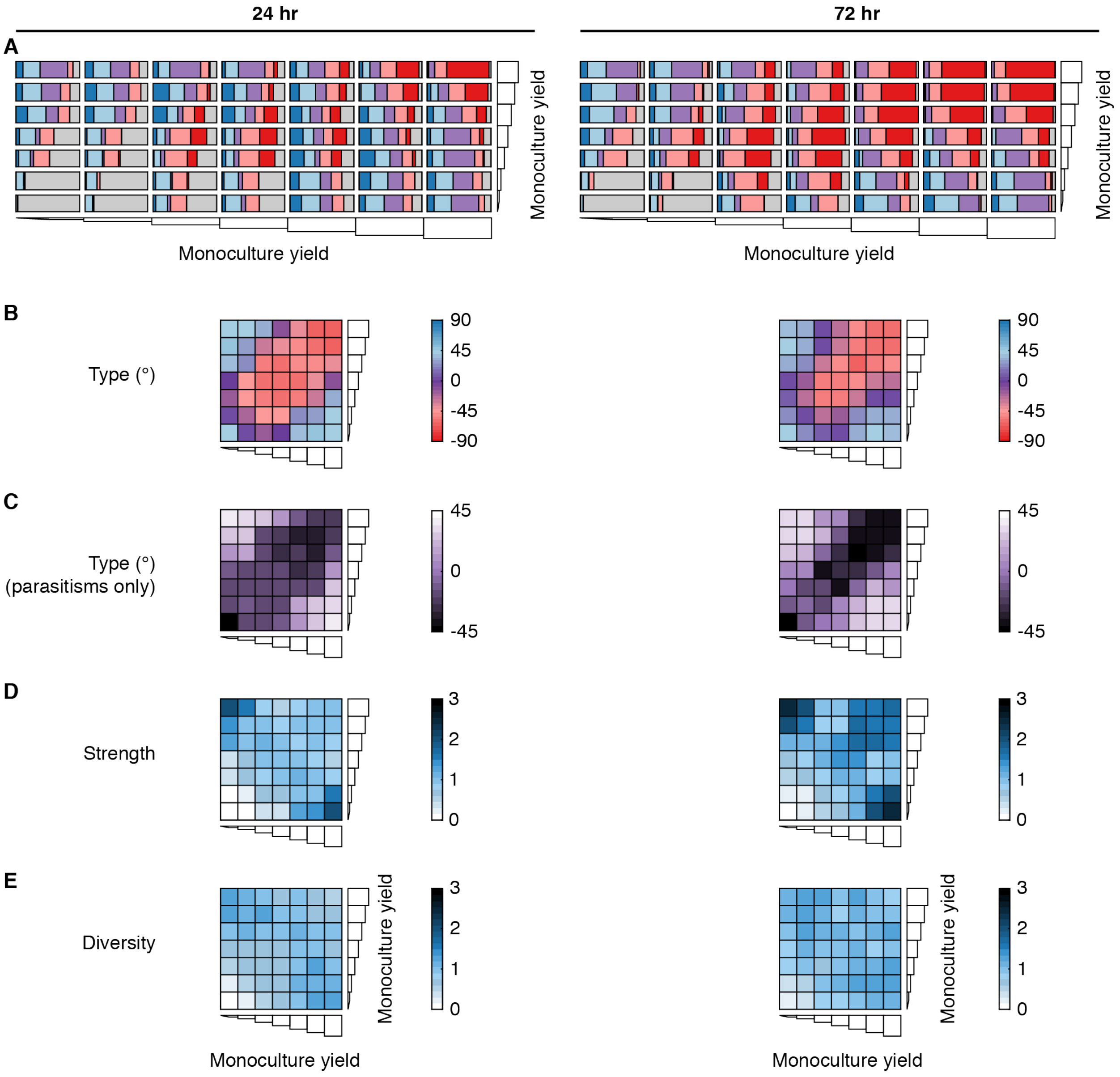


**Fig S15. Interactions by monoculture yields. A.** Interaction classification for all monoculture yield combinations. Monoculture yields were normalized based on monoculture yields at the given time point. Colors indicate interaction classification (with color legend in Fig 2). **B-D.** Average interaction metrics for all monoculture yield combinations.

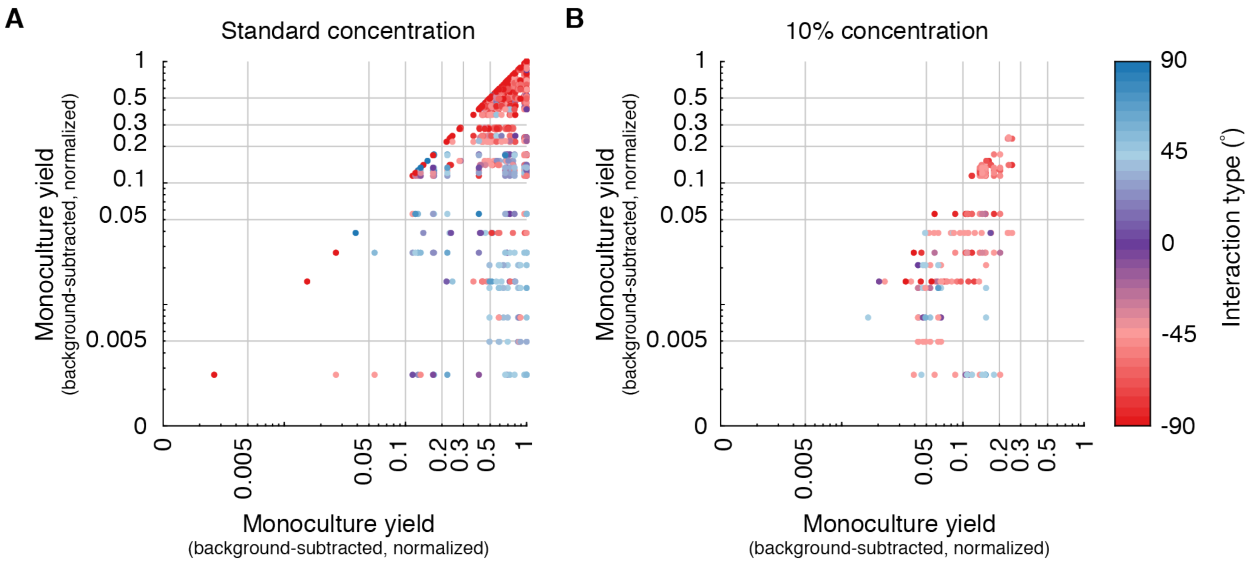


**Fig S16. Interactions arranged by monoculture yields for standard and 10% concentration.** At standard concentration (0.5% w/v) (left) and 10% concentration (0.05% w/v) (right) of carbon sources, monoculture yields and interactions were both different. All interactions at 72 hr.

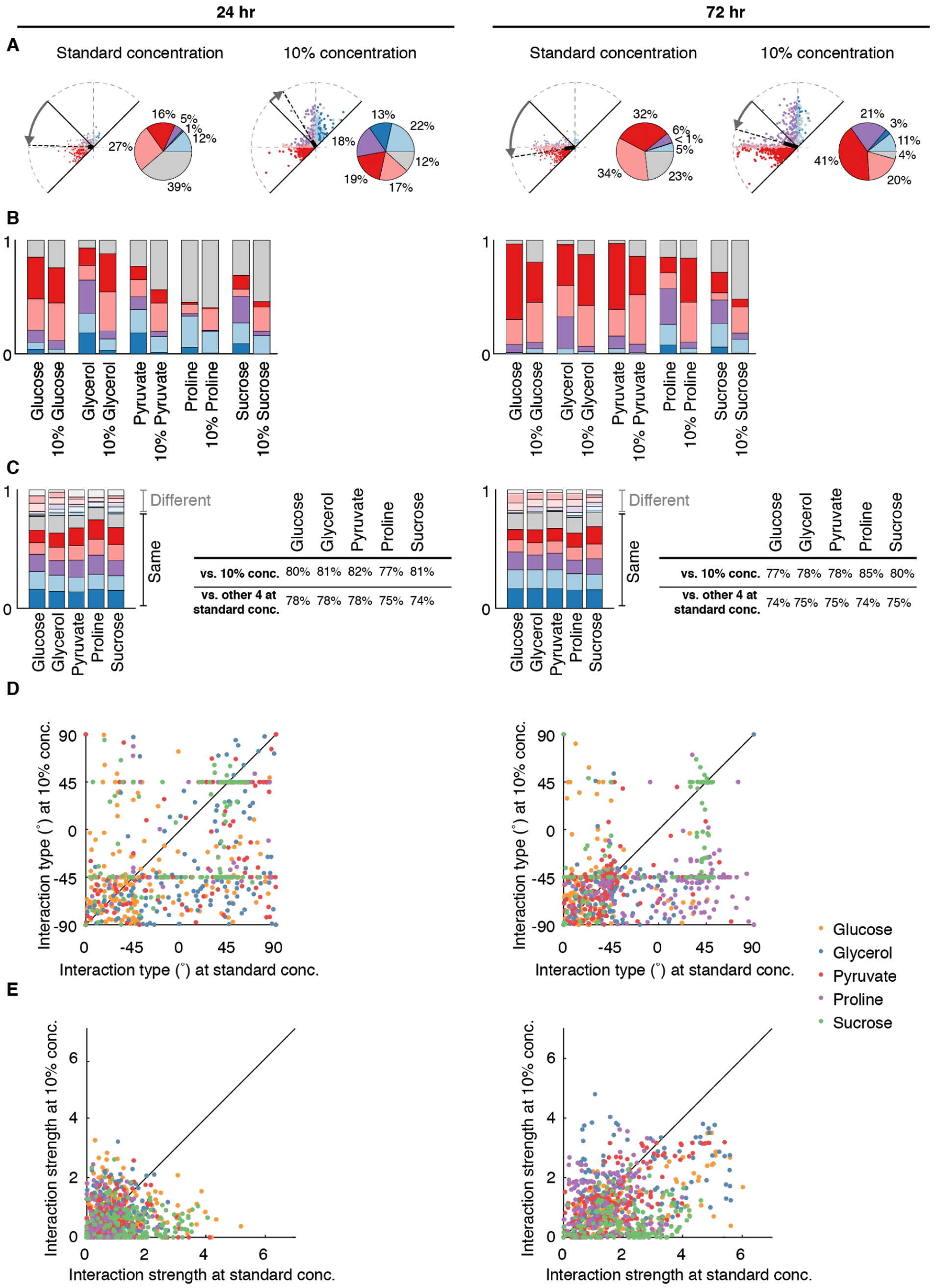


**Fig S17. Interactions with carbon sources at lower concentration. A.** All interactions occurring on 5 carbon sources (glucose, glycerol, pyruvate, proline, and sucrose) at standard concentration (0.5% w/v) and 10% concentration (0.05% w/v). **B.** Interaction classifications per carbon sources for standard and 10% concentrations. Colors indicate interaction classification (with color legend in Fig 2) **C.** Interaction classifications that are the same between the standard and 10% conditions (unmuted colors) and different (muted colors). The tables indicate the percentage agreement in interaction classification between the standard and 10% concentrations (first row) and the average percentage agreement with the 4 other carbon sources at standard concentration (second row). **D-E.** The interaction type and strength for all strain pairs at standard vs. 10% concentration, colored by carbon source.

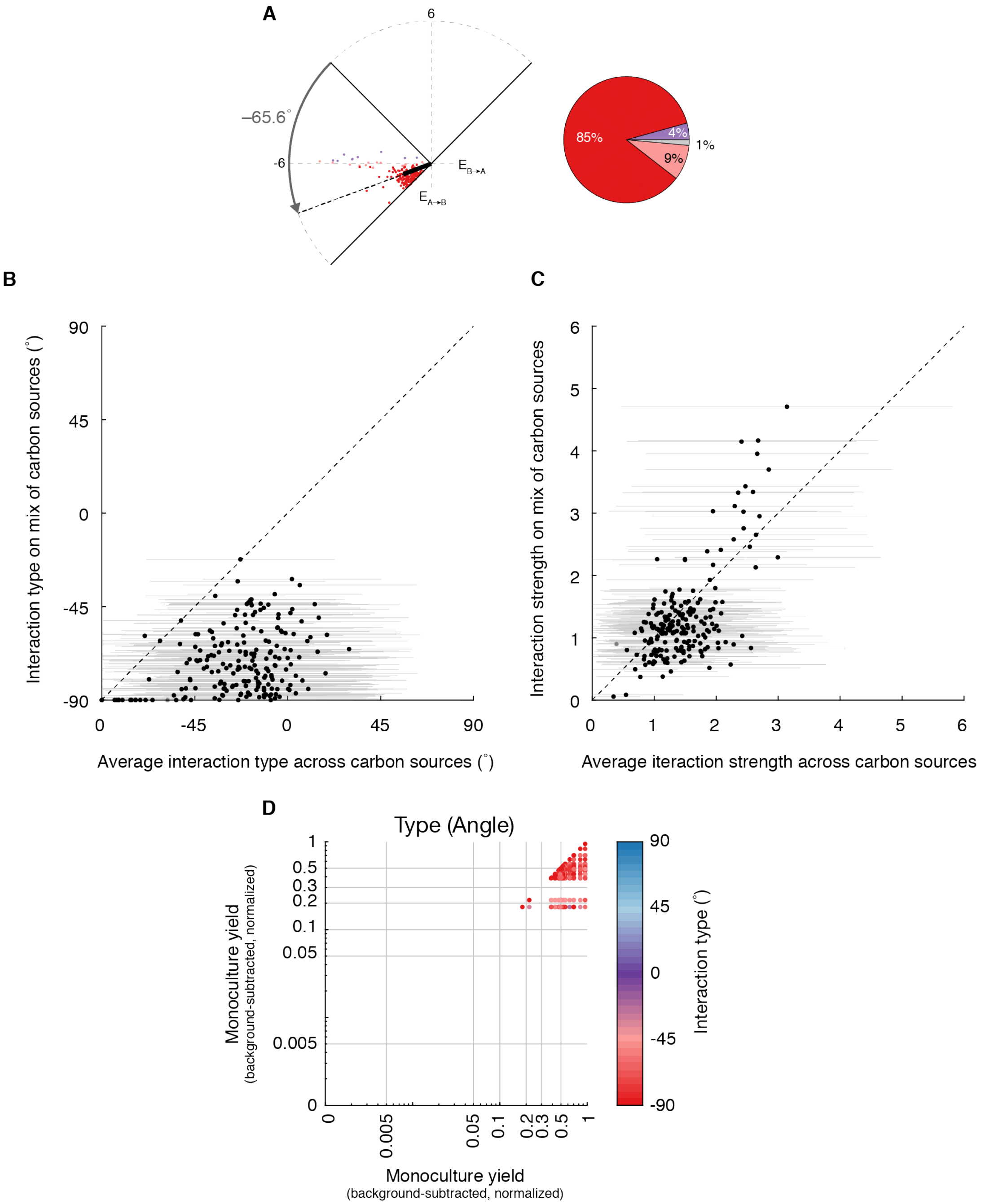


**Fig S18. Interactions on a mix of carbon sources. A.** All interactions on an even mix of 33 carbon sources (each at 0.015% w/v, total concentration 0.5% w/v). Colors indicate interaction classification (with color legend in Fig 2). **B.** Interactions arranged by monoculture yield. **C.** Comparison of interaction type for each coculture on the mix of carbon sources and mean interaction type on all 33 single carbon sources. Error bars indicate standard deviation. **D.** Comparison of interaction strength for each coculture on the mix of carbon sources and Mean interaction strength on all 33 single carbon sources. Error bars indicate standard deviation. All interactions at 72 hr.

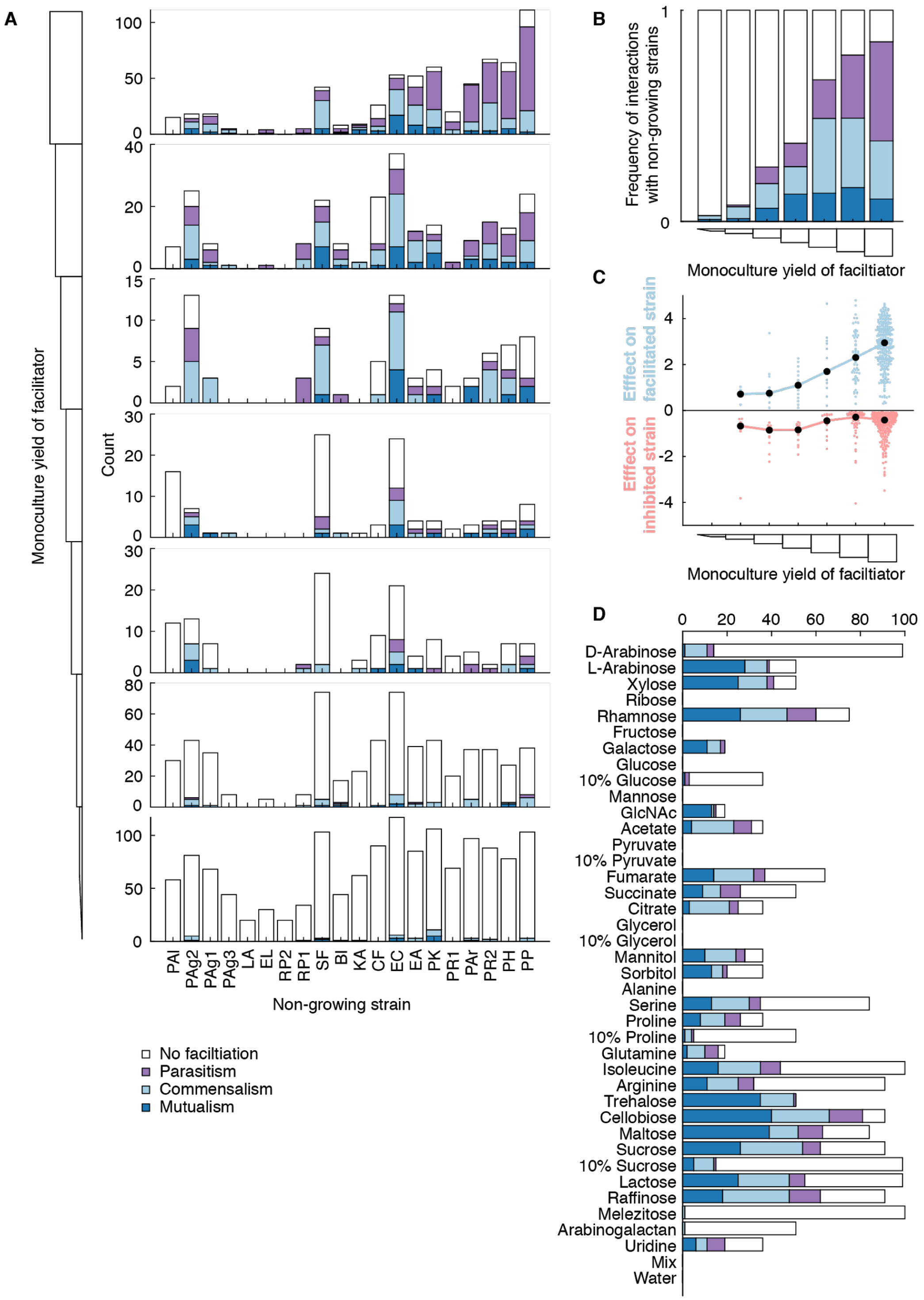


**Fig S19. Interactions containing obligate facilitation. A.** Interactions between growers and non-growers separated by monoculture yield of the facilitator. **B.** Each bar represents all interactions for each subplot in panel A (normalized to 1). **C.** Effect sizes for parasitisms where one strain was obligately facilitated. **D.** Interactions between growers and non-growers separated by carbon source.

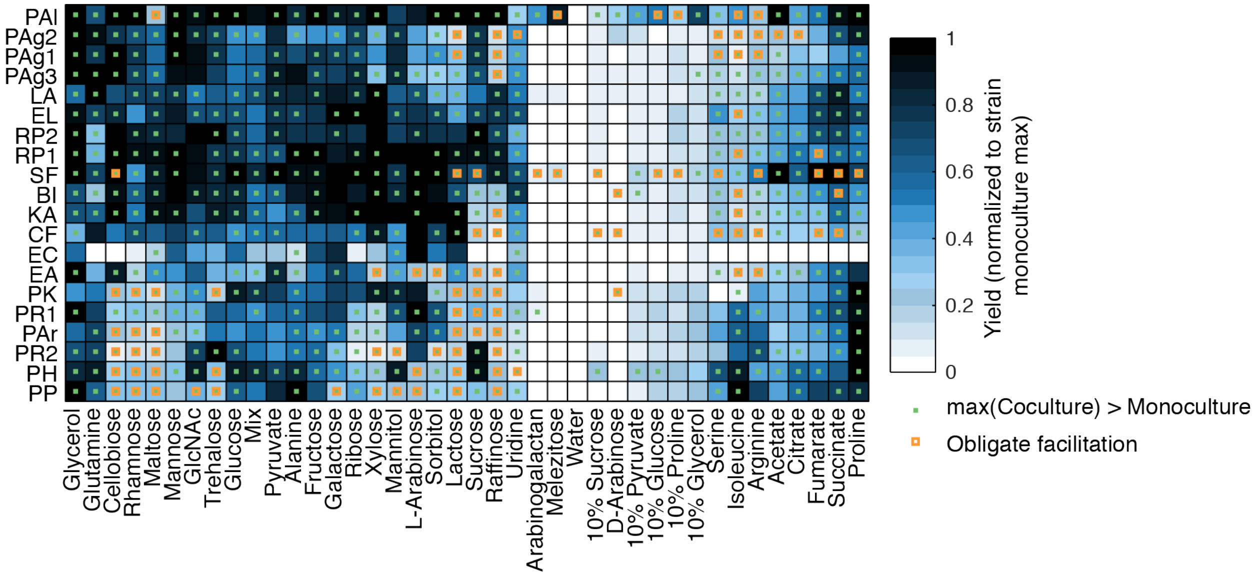


**Fig S20. Maximum yields across all dataset.** Yield values (median GFP measurement across replicates) were background-subtracted (background = median yield of labeled strain with no carbon) and normalized to the maximum yield value observed for the given strain in monoculture. Carbon sources and strains were hierarchically clustered at the 72 hr time point based on their monoculture yields only. Green dots indicate that the maximum yield of the strain/carbon source combination occurred in a coculture was at least 0.1 normalized units greater than the monoculture yield. Orange squares indicate obligate facilitation where the strain did not grow detectably on the carbon source. All interactions at 72 hr.

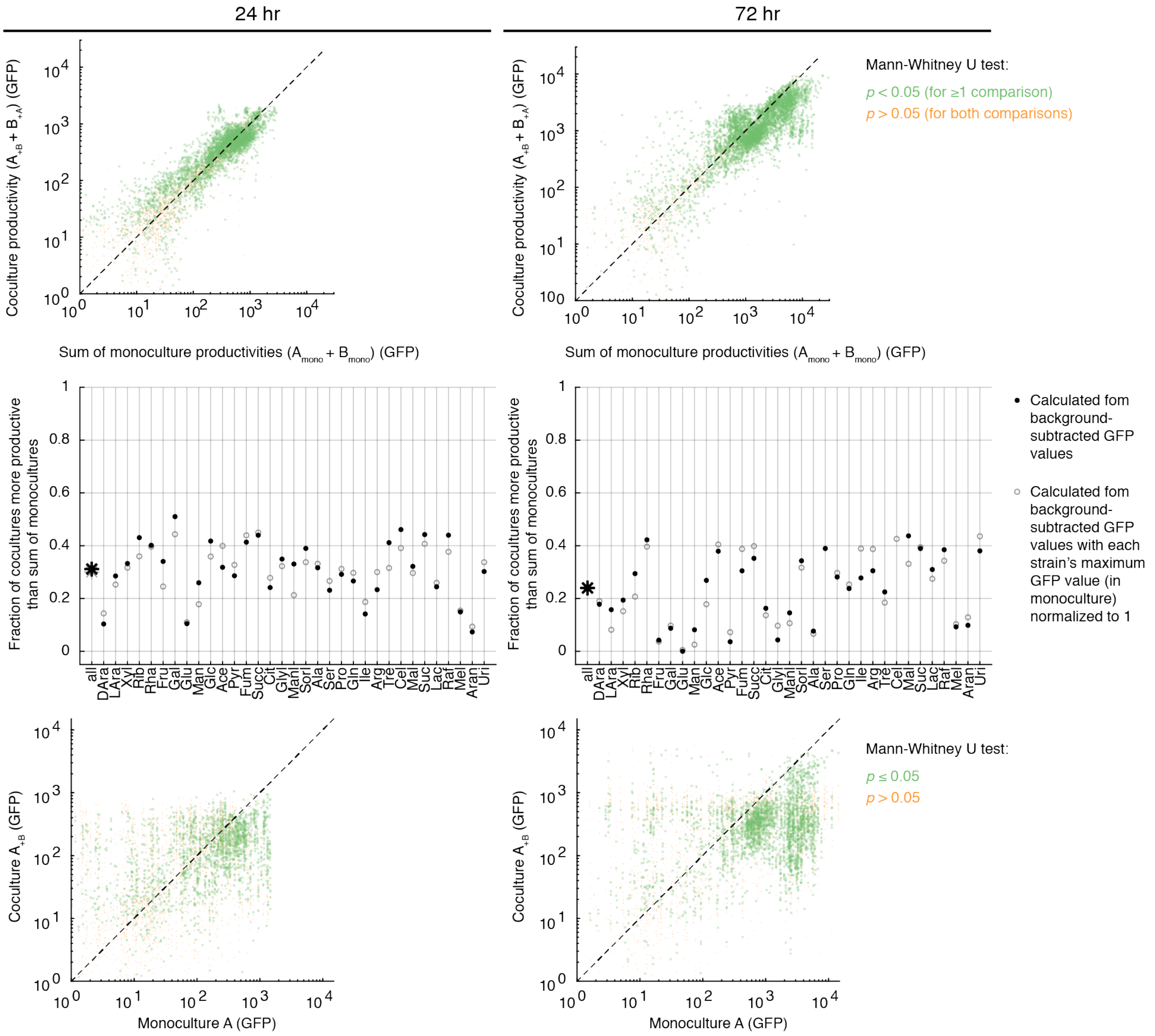


**Fig S21. Coculture vs. monoculture comparison.** (Top) Coculture productivity vs. sum of monoculture productivities. Each point represents a coculture/carbon source combination. The 33 0.5% w/v carbon sources are included. Yields calculated as background-subtracted GFP. (Middle) The fraction of cocultures more productive than the sum of monocultures for total dataset (star) and broken down by each carbon source. The solid black circles represent fractions calculated based on background-subtracted GFP values. The open gray circles represent values calculated based on background-subtracted GFP values that were then normalized to the maximum monoculture value for each strain (as in **Fig S2**). (Bottom) The yield in coculture vs. monoculture. Each point represents a coculture/carbon source combination. Yield calculated as background-subtracted GFP.

| No. | Source | Order | Closest Seqmatch (16S Gene) | Strain shorthand | Notes |
|---|---|---|---|---|---|
| 1 | Middlesex Fells | *Enterobacterales* | *Ewingella americana* | EA | |
| 2 | Middlesex Fells | *Enterobacterales* | *Raoultella planticola* | RP1 | |
| 3 | Middlesex Fells | *Enterobacterales* | *Buttiauxella izardii* | BI | |
| 4 | Middlesex Fells | *Enterobacterales* | *Citrobacter freundii* | CF | |
| 5 | Killian Court | *Enterobacterales* | *Pantoea agglomerans* | PAg1 | |
| 6 | Killian Court | *Enterobacterales* | *Klebsiella aerogenes* | KA | |
| 7 | Killian Court | *Enterobacterales* | *Raoultella planticola* | RP2 | |
| 8 | Killian Court | *Enterobacterales* | *Pantoea agglomerans* | PAg2 | |
| 9 | Killian Court | *Enterobacterales* | *Serratia fonticola* | SF1 | |
| 10 | Killian Court | *Enterobacterales* | *Lelliottia amnigena* | LA | |
| 11 | Killian Court | *Enterobacterales* | *Pantoea allii* | PAl | |
| 12 | Killian Court | *Enterobacterales* | *Pantoea agglomerans* | PAg3 | |
| 13 | Harvard Yard | *Pseudomonadales* | *Pseudomonas helmanticensis* | PH | |
| 14 | Lalicata Compost | *Pseudomonadales* | *Pseudomonas rhodesiae* | PR1 | |
| 15 | Tech Square Yard | *Pseudomonadales* | *Pseudomonas plecoglossicida* | PP | |
| 16 | Office Plant | *Enterobacterales* | *Enterobacter ludwigii* | EL | |
| 17 | Charles River Soil | *Pseudomonadales* | *Pseudomonas rhodesiae* | PR2 | |
| 18 | Charles River Soil | *Pseudomonadales* | *Pseudomonas koreensis* | PK | |
| 19 | Harvard Yard | *Pseudomonadales* | *Pseudomonas arsenicoxydans* | PAr | Needed to be repeated. |
| 20 | Lab strain | *Enterobacterales* | *Escherichia coli* | EC | Needed to be repeated. |
| | Killian Court | | *Serratia fonticola* | SF2 | Excluded from analysis. |
| | *C. elegans* gut | | *Citrobacter braakii* | CB | Excluded from analysis. |

**Table S1. Strains used in coculture experiment.**

| No. | Isolate ID | Closest Seqmatch (16S Gene) | Included in Screen | Source | Isolation Media |
|---|---|---|---|---|---|
| 1 | M-C10 | *Buttiauxella izardii* | Yes | Middlesex Fells | Brucella |
| 2 | M-C12 | *Citrobacter braakii* | | Middlesex Fells | Campylobacter |
| 3 | Cr-75 | *Citrobacter braakii* | Yes | *C. elegans* gut | TSB |
| 4 | M-G11 | *Citrobacter freundii* | Yes | Middlesex Fells | Campylobacter |
| 5 | O-G1 | *Enterobacter ludwigii* | Yes | Office Plant | Actinomycetes |
| 6 | M-B10 | *Ewingella americana* | | Middlesex Fells | pH4-M9+Glc .005%+AA .002% |
| 7 | M-A12 | *Ewingella americana* | Yes | Middlesex Fells | ATCC Medium 111 |
| 8 | M-C11 | *Ewingella americana* | | Middlesex Fells | pH4-M9+Glc .005%+AA .002% |
| 9 | Cr-24 | *Ewingella americana* | | *C. elegans* gut | TSB |
| 10 | K-A7 | *Klebsiella aerogenes* | Yes | Killian Court | M9+glc .005%+casAA .002% |
| 11 | Cr-61 | Unclassified *Leclercia* | | *C. elegans* gut | TSB |
| 12 | K-D8 | *Lelliottia amnigena* | Yes | Killian Court | Streptomyces |
| 13 | K-C10 | *Pantoea agglomerans* | Yes | Killian Court | pH5-M9+Glc .5%+AA .2% |
| 14 | K-A4 | *Pantoea agglomerans* | Yes | Killian Court | M9 + glucose .5% |
| 15 | K-F8 | *Pantoea agglomerans* | Yes | Killian Court | Bordatella-Blood Agar |
| 16 | K-G6 | *Pantoea agglomerans* | | Killian Court | Nutrient Agar - 1% |
| 17 | K-A5 | *Pantoea agglomerans* | | Killian Court | M9+glc .005%+casAA .002% |
| 18 | K-D12 | *Pantoea agglomerans* | | Killian Court | Campylobacter |
| 19 | K-E11 | *Pantoea agglomerans* | | Killian Court | ATCC Medium 111 |
| 20 | K-A12 | *Pantoea agglomerans* | | Killian Court | Bordatella-Blood Agar |
| 21 | K-F4 | *Pantoea allii* | Yes | Killian Court | LB |
| 22 | K-B3 | *Pantoea ananatis* | | Killian Court | M9 + glucose .5% |
| 23 | O-A1 | *Pseudomonas abietaniphila* | | Office Plant | TSB |
| 24 | H-E5 | *Pseudomonas arsenicoxydans* | Yes | Harvard Yard | LB |
| 25 | H-F5 | *Pseudomonas arsenicoxydans* | | Harvard Yard | Nutrient Agar |
| 26 | Rh-C8 | *Pseudomonas baetica* | | Rocky Hill Compost | ATCC Medium 111 |
| 27 | K-A8 | *Pseudomonas donghuensis* | | Killian Court | M9 + glucose .005% |
| 28 | H-D3 | *Pseudomonas helmanticensis* | Yes | Harvard Yard | TSB |
| 29 | R-D5 | *Pseudomonas koreensis* | Yes | Charles River Soil | M9 + glucose .005% |
| 30 | K-H1 | *Pseudomonas koreensis* | | Killian Court | Nutrient Agar |
| 31 | H-E3 | *Pseudomonas koreensis* | | Harvard Yard | Nutrient Agar |
| 32 | K-H1 | *Pseudomonas koreensis* | | Killian Court | Nutrient Agar |
| 33 | O-E8 | *Pseudomonas migulae* | | Office Plant | Streptomyces |
| 34 | O-D6 | *Pseudomonas monteilii* | | Office Plant | M9+glc .005%+casAA .002% |
| 35 | T-G7 | *Pseudomonas plecoglossicida* | Yes | Tech Square Yard | ATCC Medium 111 |
| 36 | L-C5 | *Pseudomonas rhodesiae* | Yes | Lalicata Compost | LB |
| 37 | R-C1 | *Pseudomonas rhodesiae* | Yes | Charles River Soil | Nutrient Agar |
| 38 | L-A6 | *Pseudomonas rhodesiae* | | Lalicata Compost | Streptomyces |

| | | | | | |
|---|---|---|---|---|---|
| 39 | R-C6 | *Pseudomonas rhodesiae* | | Charles River Soil | Campylobacter |
| 40 | H-A7 | *Pseudomonas silesiensis* | | Harvard Yard | Nutrient Agar |
| 41 | H-A1 | *Pseudomonas vancouverensis* | | Harvard Yard | LB |
| 42 | H-G1 | *Pseudomonas vancouverensis* | | Harvard Yard | LB |
| 43 | K-H12 | *Raoultella planticola* | | Killian Court | pH4-M9+Glc .005%+AA .002% |
| 44 | M-B5 | *Raoultella planticola* | Yes | Middlesex Fells | M9 + glucose .005% |
| 45 | K-A10 | *Raoultella planticola* | Yes | Killian Court | Streptomyces |
| 46 | Cr-77 | *Raoultella planticola* | | *C. elegans* gut | TSB |
| 47 | Cr-86 | *Raoultella planticola* | | *C. elegans* gut | TSB |
| 48 | K-H8 | *Serratia fonticola* | | Killian Court | TSB |
| 49 | K-D4 | *Serratia fonticola* | Yes | Killian Court | Actinomycetes |
| 50 | K-H10 | *Serratia fonticola* | Yes | Killian Court | Streptomyces |

**Table S2. Bacterial isolates fluorescently labeled with sGFP2 (pMRE132).**

| No. | Biochemical class | Full compound | Name used | Abbreviation | Final Conc. (% w/v) |
|---|---|---|---|---|---|
| 1 | Monosaccharide | D-Arabinose | D-Arabinose | DAra | 0.5 |
| 2 | Monosaccharide | L-Arabinose | L-Arabinose | LAra | 0.5 |
| 3 | Monosaccharide | D-Xylose | Xylose | Xyl | 0.5 |
| 4 | Monosaccharide | D-Ribose | Ribose | Rib | 0.5 |
| 5 | Monosaccharide | L-Rhamnose monohydrate | Rhamnose | Rha | 0.5 |
| 6 | Monosaccharide | D-Fructose | Fructose | Fru | 0.5 |
| 7 | Monosaccharide | D-Galactose | Galactose | Gal | 0.5 |
| 8 | Monosaccharide | D-Glucose | Glucose | Glu | 0.5 |
| 9 | Low conc. | 0.1X D-Glucose | Glucose01 | Glu01 | 0.05 |
| 10 | Monosaccharide | D-Mannose | Mannose | Man | 0.5 |
| 11 | Monosaccharide | N-Acetyl-D-glucosamine | GlcNAc | Glc | 0.5 |
| 12 | TCA | Sodium acetate | Acetate | Ace | 0.5 |
| 13 | TCA | Sodium pyruvate | Pyruvate | Pyr | 0.5 |
| 14 | Low conc. | 0.1X Sodium pyruvate | Pyruvate01 | Pyr01 | 0.5 |
| 15 | TCA | Sodium fumarate dibasic | Fumarate | Fum | 0.5 |
| 16 | TCA | Disodium succinate | Succinate | Succ | 0.5 |
| 17 | TCA | Sodium citrate dihydrate | Citrate | Cit | 0.5 |
| 18 | Sugar alcohol | Glycerol | Glycerol | Glyl | 0.5 |
| 19 | Low conc. | 0.1X Glycerol | Glycerol01 | Glyl01 | 0.05 |
| 20 | Sugar alcohol | D-Mannitol | Mannitol | Manl | 0.5 |
| 21 | Sugar alcohol | D-Sorbitol | Sorbitol | Sorl | 0.5 |
| 22 | Amino acid | L-Alanine | Alanine | Ala | 0.5 |
| 23 | Amino acid | L-Serine | Serine | Ser | 0.5 |
| 24 | Amino acid | L-Proline | Proline | Pro | 0.5 |
| 25 | Low conc. | 0.1X L-Proline | Proline01 | Pro01 | 0.05 |
| 26 | Amino acid | L-Glutamine | Glutamine | Gln | 0.5 |
| 27 | Amino acid | L-Isoleucine | Isoleucine | Ile | 0.5 |
| 28 | Amino acid | L-Arginine | Arginine | Arg | 0.5 |
| 29 | Disaccharide | D-Trehalose dihydrate | Trehalose | Tre | 0.5 |
| 30 | Disaccharide | D-Cellobiose | Cellobiose | Cel | 0.5 |
| 31 | Disaccharide | D-Maltose monohydrate | Maltose | Mal | 0.5 |
| 32 | Disaccharide | D-Sucrose | Sucrose | Suc | 0.5 |
| 33 | Low conc. | 0.1X D-Sucrose | Sucrose01 | Suc01 | 0.05 |
| 34 | Disaccharide | D-Lactose monohydrate | Lactose | Lac | 0.5 |
| 35 | Trisaccharide | D-Raffinose pentahydrate | Raffinose | Raf | 0.5 |
| 36 | Trisaccharide | D-Melezitose monohydrate | Melezitose | Mel | 0.5 |
| 37 | Arabinogalactan | Arabinogalactan | Arabinogalactan | Aran | 0.5 |
| 38 | Uridine | Uridine | Uridine | Uri | 0.5 |
| 39 | Mix | Mix | Mix | Mix | 0.5 |
| 40 | Water | Water | Water | Wat | – |

**Table S3. Carbon sources used in coculture experiment.**
